# Sampling Conformational Ensembles of Highly Dynamic Proteins via Generative Deep Learning

**DOI:** 10.1101/2024.05.05.592587

**Authors:** Talant Ruzmetov, Ta I Hung, Saisri Padmaja Jonnalagedda, Si-han Chen, Parisa Fasihianifard, Zhefeng Guo, Bir Bhanu, Chia-en A. Chang

## Abstract

Proteins are inherently dynamic, and their conformational ensembles are functionally important in biology. Large-scale motions may govern protein structure–function relationship, and numerous transient but stable conformations of Intrinsically Disordered Proteins (IDPs) can play a crucial role in biological function. Investigating conformational ensembles to understand regulations and disease-related aggregations of IDPs is challenging both experimentally and computationally. In this paper we first introduce a deep learning-based model, termed Internal Coordinate Net (ICoN), which learns the physical principles of conformational changes from Molecular Dynamics (MD) simulation data. Second, we selected interpolating data points in the learned latent space that rapidly identify novel synthetic conformations with sophisticated and large-scale sidechains and backbone arrangements. Third, with the highly dynamic amyloid-β_1-42_ (Aβ42) monomer, our deep learning model provided a comprehensive sampling of Aβ42’s conformational landscape. Analysis of these synthetic conformations revealed conformational clusters that can be used to rationalize experimental findings. Additionally, the method can identify novel conformations with important interactions in atomistic details that are not included in the training data. New synthetic conformations showed distinct sidechain rearrangements that are probed by our EPR and amino acid substitution studies. This approach is highly transferable and can be used for any available data for training. The work also demonstrated the ability of deep learning to utilize learned natural atomistic motions in protein conformation sampling.

## Introduction

Proteins are complex and have dynamic properties. Conformational ensembles of proteins are essential for performing biological processes, including enzyme activities, protein folding, protein–ligand binding, and protein aggregation ^1,2^. Characterization of protein conformations allows researchers to understand protein function, activity, and mechanisms. However, the tasks may be daunting and are even more challenging for investigating intrinsically disordered proteins (IDPs), for which numerous conformations can be functionally important. Some IDPs, such as amyloid-β_1-42_ (Aβ42), are related to several protein aggregation-related diseases, including Alzheimer’s disease ^3^. Revealing important conformations with atomistic details is crucial to fully understand the function of a protein and its regulation involved in basic science, therapeutics design and biotechnology.

Experimental techniques determine protein structures and probe their dynamics, and bioinformatics tools such as AlphaFold2 and RoseTTAFold can accurately predict structures from given protein sequences ^4–6^. However, these techniques typically provide a handful of structures with limited information on dynamics. Advanced computational methods are widely used to investigate protein dynamics and sample protein conformational ensembles. A powerful method is all-atom molecular dynamic (MD) simulations, which are based on physical principles to sample protein conformations and model their dynamics ^7,8^. However, many stable protein conformations are separated by free energy barriers: an MD run can take excessively long computational time (i.e., beyond the microsecond timescale) to cross energy barriers for sampling various conformations. Other methods developed to tackle sampling limitations include conformational search methods, MD-based enhanced sampling techniques, and Monte Carlo simulations ^6,9–12^. A popular analysis tool, Principal Component Analysis (PCA), has been used to extract information from sampled protein conformations to guide conformational search ^13,14^. Low-mode-based conformational search methods use Normal Mode Analysis (NMA) to perturb given structures along their low-mode eigenvectors to generate new conformations ^15–18^.

Although these sampling approaches effectively extend the time scale of MD simulations, efficiently obtaining adequate conformations for highly flexible protein systems and/or overcoming significant energy barriers is still challenging.

Artificial Intelligence (AI) provides an alternative approach for accelerating the generation of protein conformational ensembles. Novel machine learning/deep learning (ML/DL) approaches analyze simulation results to further guide conformational search with enhanced sampling techniques ^19–28^. ML/DL optimizes coarse-grain models to speed up conformation transitions with preserved atomistic interactions ^29,30^. Several DL models accelerate protein conformational searches, such as training neural networks to model the distribution of conformational ensembles with generative modeling ^31–40^. By training with conformations obtained from MD simulations, these approaches can quickly generate conformational ensembles and bypass the kinetic barriers, the bottleneck of MD simulation.

Generative models study conformations of chemical compounds and proteins in various settings; examples include coarse-grained models and protein conformations presented by a backbone ^29,41^. Ideally, the generative models can find new and thermodynamically stable protein conformational ensembles not seen in the training dataset obtained from MD simulations, as demonstrated in recent publications focusing on sampling conformations for IDPs ^42–47^.

Nevertheless, proteins have complex conformational ensembles, and many of these transitions are driven by backbone motions and, more importantly, sidechain interactions such as salt bridges, hydrogen bonds, and hydrophobicity. As a result, to find new protein conformations, features that can accurately describe all possible motions in atomistic details are critical in generative AI models.

In this study, we present a novel technique with improved input representation, and a generative sampling method using an autoencoder-based backbone, termed Internal Coordinate Net (ICoN). Our proposed pipeline is trained on datasets from MD simulations to identify new conformations efficiently and accurately for highly flexible protein systems. ICoN uses novel features and an atomistic bond-angle-torsion (BAT)-based vector representation, vBAT, to smoothly rotate various dihedral angles that lead to new conformations. Our model learns the essential dihedral motions which follow the physical principles that govern molecular motions, and the 3D latent space contains transformations of various degrees of freedom (DOF), mainly dihedral rotations, to allow for efficient conformational searches. Essential protein motions analyzed by dihedral Principal Component Analysis (dPCA) are clearly revealed in the latent space, and generating new conformations is achieved by data-point interpolation within the latent space. Using a fraction (1%) of MD data as a training set, we demonstrated that ICoN rapidly found thousands of new, conformationally distinct, and thermodynamically stable conformations for αB-crystallin57-69 and the Aβ42 monomer (**Figure 1**) in a few minutes using a single gaming GPU card. In addition to Aβ42 monomer, we also used αB-crystallin 57–69, a disease-related long-lived peptide with D-amino acid and two serine isomerization sites and a potential therapeutic target, to demonstrate molecular motions revealed in the latent space ^48^. Post-analysis of our novel synthetic Aβ42 monomer conformations revealed various new conformations, including new Arg5-Ala42 contacts and a new Asp23-Lys28 salt bridge not seen in the MD training set but reported in existing publications. Our synthetic conformations covered reported structures that were suggested to be less toxic or more prone to oligomerization ^49–55^.

**Figure 1.**
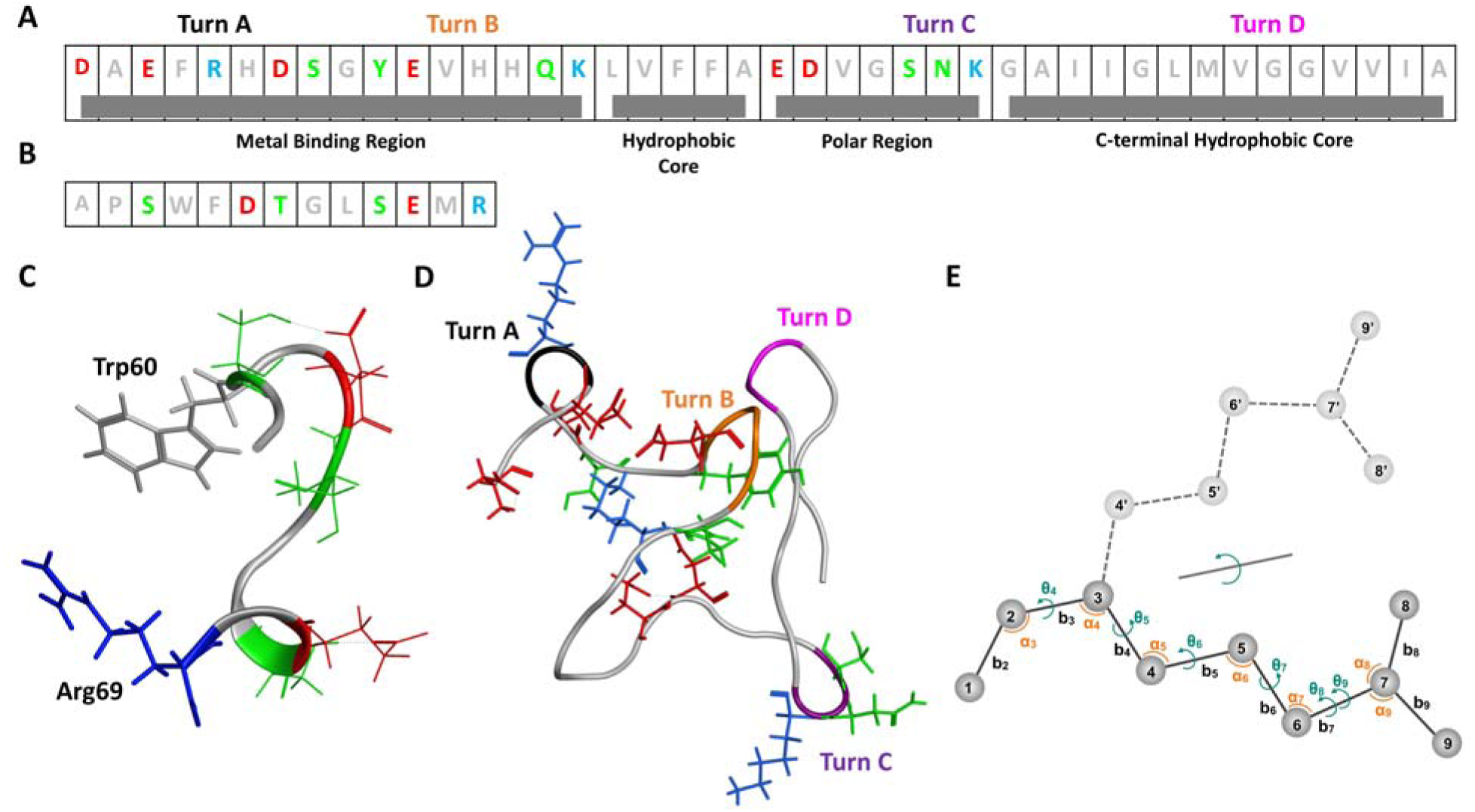
(A) Sequence of 42-residue amyloid beta peptide (Aβ42). (B) Sequence of 13-residue αB-crystallin57-69. (C) A synthetic conformation of αB-crystallin57-69 with intramolecular charge and aromatic ring interactions between Trp60 and Arg69. Red: negative charge residues; Blue: positive charge residues; Green: polar; Grey: neutral and non-polar residues. (D) A as illustrated in Figure 6B synthetic conformation of Aβ42 with all 4 turns A-D. (E) Classical internal bond-angle-torsion (BAT) coordinate representation (Z-matrix) for a small molecule. Atoms 1−3 are termed root atoms for this molecule. Atoms i > 3 are defined by (bi, ai, θi), where b, a and θ are bond, angle and torsions, respectively. For example, atom 4 is presented by (b4, a4, θ4). The dashed light gray line shows the smooth rotation of the bond between atoms 2 and 3 (θ4).

Our model provides a computationally efficient approach to identify new conformations for highly flexible protein systems. The novel synthetic Aβ42 monomer conformations also provide insights into Aβ42 oligomerization.

## Results

### ICoN model architecture, training, and validation

#### Molecular representation using the vBAT coordinate

The ICoN model is a latent space-guided generative sampling using an autoencoder model trained on protein conformational ensembles to learn physical properties that govern conformational transitions in MD simulations (**Figure 2**).

**Figure 2.**
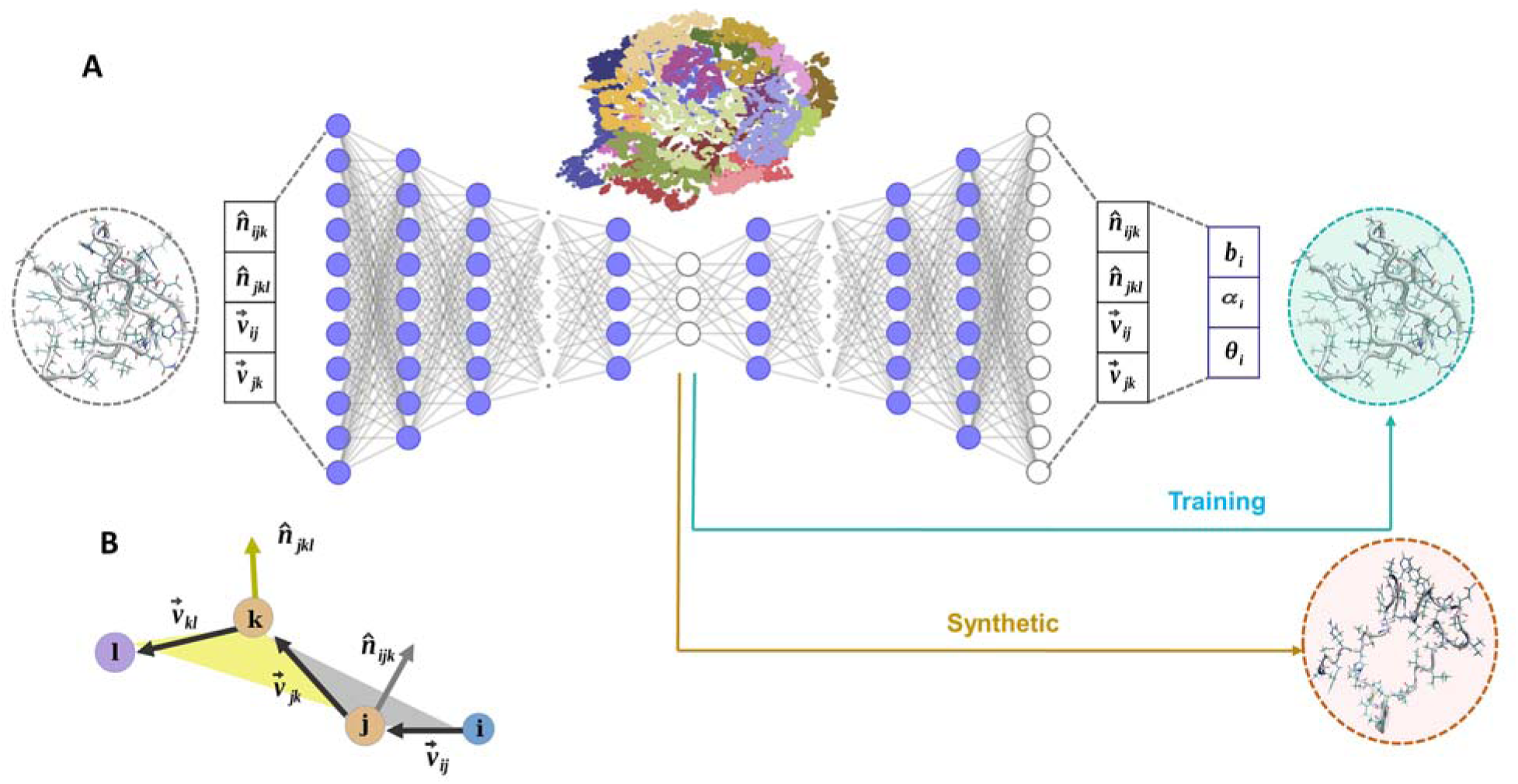
Overview of the Internal Coordinate Net (ICoN) network architecture. (A) The input data are thousands of protein conformations represented by BAT-based vBAT features. The molecular conformations have 3N-6 degrees of freedom and 4×3(N-3) dimension features, where N is the number of atoms. The high dimensions are compressed by a series of fully connected layers into a 3D latent space using an encoder. The entire process is then inverted back to the original dimensions using a decoder and then converted back to Cartesian coordinates. (B) An illustration of vBAT features using 4 atoms with indexes, i, j, k, and l. The vectors connecting the atoms 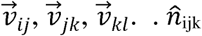 and 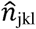 are unit vectors normalized to the planes constructed by atoms i-j-k and j-k-l, respectively.

Because the dihedral rotations are the major motions that determine the conformations, using BAT to accurately describe the concerted motions of multiple dihedral rotations is critical.

Dihedral rotations have a periodicity issue, so we used a vector called vBAT to avoid periodicity (**Figure 2B** and details in Supplementary Material). All-atom vBAT internal coordinate representation is inherently equivariant to rotations and translations and can exactly convert vBAT vectors to Cartesian coordinates without any additional approximation. However, coordinate conversions can be time-consuming. Specifically, reconstructing Cartesian coordinate from BAT requires defining one atom at a time which is time-consuming in the network. Thus, our model implemented GPU acceleration to perform coordinate transformations; converting 5,000 frames from vBAT to Cartesian coordinates took < 0.2 s for Aβ42. **Figure S2** demonstrates the efficiency in coordinate transformations that successfully avoided a computation bottleneck with the use of internal coordinates.

#### ICoN network architecture and training

The neural network that we use for sampling is based on autoencoder architecture (**Figure 2A**). Classical BAT coordinates (the Z matrix) have 3N-6 DOF for presenting the internal motions of a molecule, where N is the number of atoms, and the 6 external translation and rotation DOF are eliminated. Because vBAT uses the vectors instead of one value for each bond, angle and dihedral DOF, we have 4×3×(N-3) features for each molecule. As a result, αB-crystallin57-69 and Aβ42 have 2376 and 7488 features, respectively. The hyperparameter training in the encoder resulted in 7 layers to reduce a large dimension representation into 3D, so each molecular conformation can be presented in the 3D latent space (**Figures 2A** and **S3**). The decoding process brought the representation from 3D back to the original dimensions with the same number of layers but in reverse order. Each conformation is converted from vBAT to Cartesian coordinates for further analysis. Throughout the iterative training process, in addition to learning a relationship between a vast number of features, the network also learns how to meaningfully compress the representation into the lower-dimensional 3D latent space by performing nonlinear dimensionality reduction.

Notably, although this study used 3D latent space for easy observation of conformation distribution and visualization of interpolation, the network can reduce 4×3×(N-3) features to any dimension desired by users. We used 3D to achieve efficient training and reduced memory consumption as well. It took 4 min with one NVIDIA 1080Ti GPU card to perform 15000 epochs for successful training with 10,000 frames of Aβ42.

#### Validation of the models by conformation reconstruction

A successfully trained model should accurately reconstruct a protein conformation from the reduced 3D representation to the original conformation. To evaluate the accuracy, we presented conformations from our validation set using the 3D representation and then reconstructed them back to atomistic Cartesian representation and computed the root mean square deviation (RMSD) of all heavy atoms between the 2 conformations, the original and reconstructed ones. A few representative conformations are illustrated in **Figure S4**: αB-crystallin57-69 has RMSD < 0.9Å and Aβ42 has RMSD < 1.3Å. As shown in **Table 1**, with our model, both backbone and sidechain conformations are reconstructed back to the original input structure with average dihedral RMSD values reported.

**Table 1.**
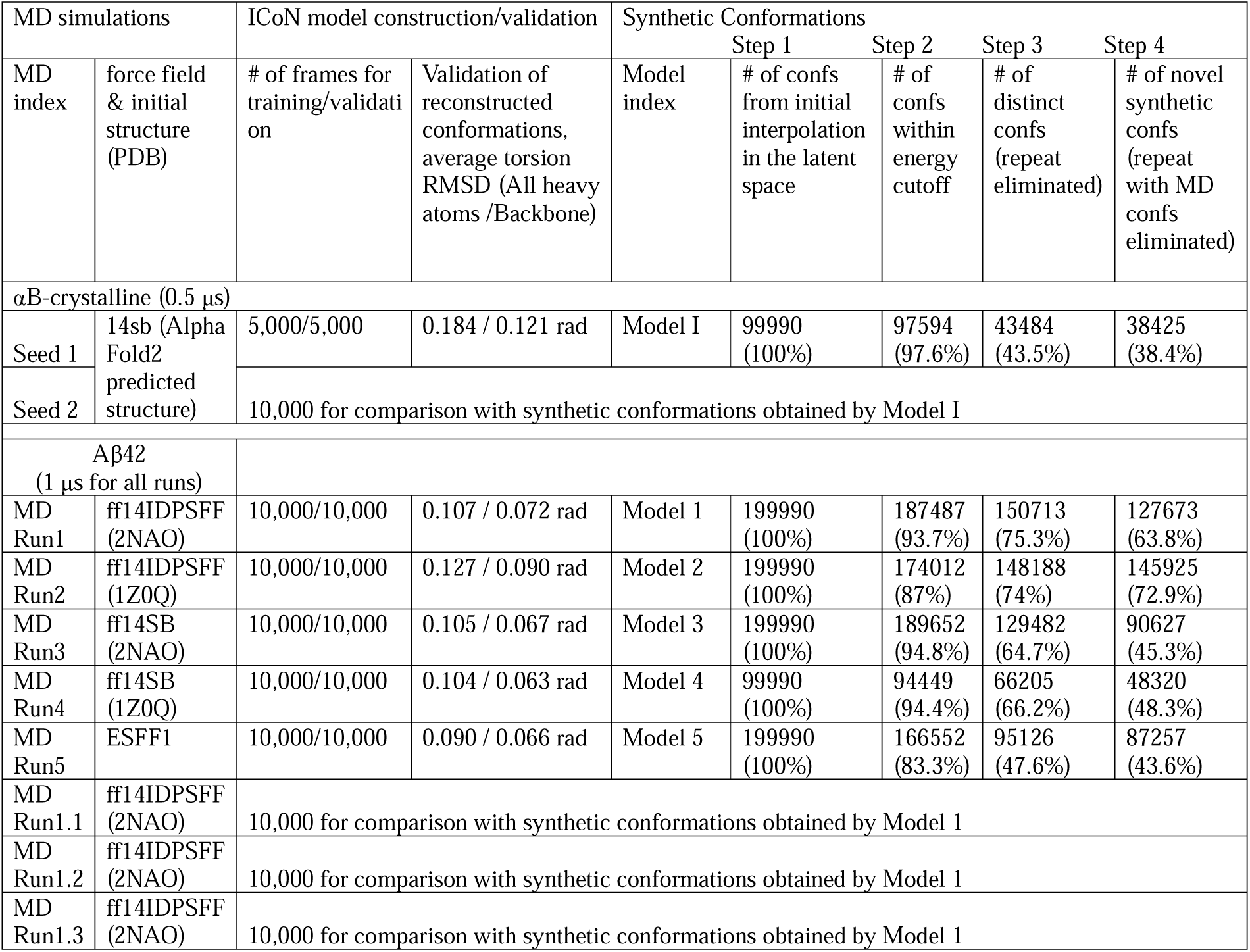
Summary of molecular dynamics (MD) simulations, numbers of MD frames used for ICoN model training and validation, and numbers of conformations generated from ICoN models in each step. The hyperparameters of Aβ42 ICoN model were obtained from dataset MD Run1, and all Models 1 to 5 use the same hyperparameters. Energy cutoff was-200 and-400 kcal/mol for αB-crystalline and Aβ42. Two conformations are treated as repeats when the computed heavy atom RMSD is smaller than 1 Å in Step 3. In Step 4, the RMSD cutoff is 1 Å and 1.2 Å for αB- crystalline and Aβ42, respectively.

We also performed detailed comparisons between the original and reconstructed conformations, such as the distribution of each dihedral rotation and their correlations, to ensure that all the properties were accurately reproduced (**Figures S5-S8**). Notably, both αB-crystallin57-69 and Aβ42 exhibit large-scale protein motions included in our training set. The validation demonstrates the robustness of our model to learn a diverse range of conformational states with high precision, and that we can describe small sidechain rotations accurately.

### Learned physics in the latent space of αB-cristallin57-69

Because our ICoN model learns important dihedral rotations that determine molecular conformations, the latent space stores information that leads to smooth conformational transitions between 2 data points, such as various sets of concerted dihedral rotations. Moreover, we can visualize conformation transitions and clusters and can perform data analysis in the 3D latent space (**Figures 3 and S10**). We demonstrated that the DL model learned the physics governing molecular motions, and the information captured in the latent space can be effectively utilized.

**Figure 3.**
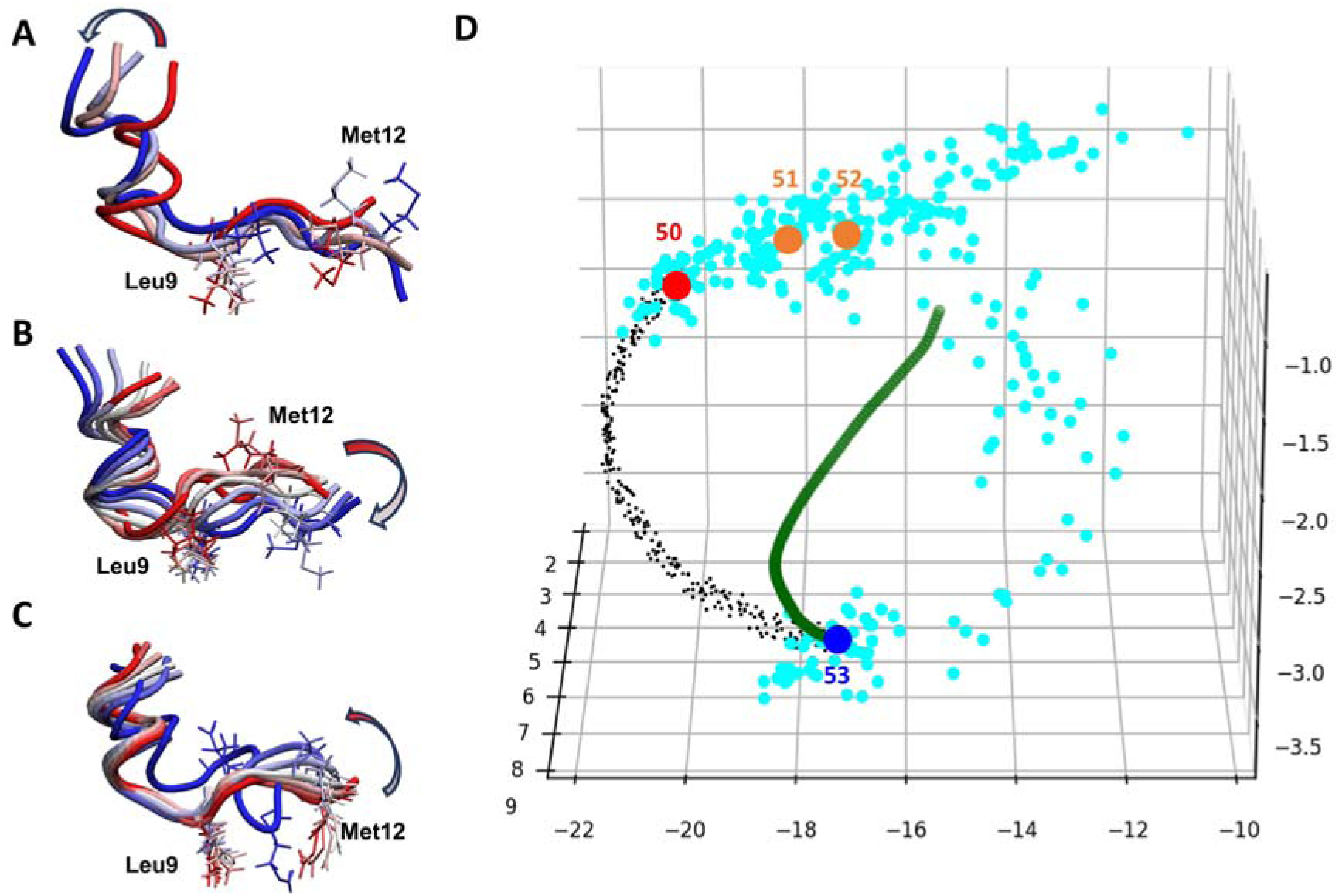
Conformational transitions of αB-crystallin57-69 peptide. (A) Super-imposed conformations used for DL training (Conf indexes #50 to #53). (B) Conformational transitions led by distortions using the first dihedral principal component (PC) mode. (C) Conformational transitions led by interpolation between training Conf indexes #50 (red dot) and #53 (dark blue dot). Leu9 and Met12, which have the most fluctuations, are presented in licorice. (D) 3D latent space from the ICoN model. Light blue dots present conformations saved every 1 ps in MD, with a total of 300 dots (300-ps simulation length). The MD frames were re-saved every 100 ps for training, and 4 frames are presented in the plot (red, blue and orange dots). The green dot line illustrates conformation distortions using the first PC mode, with the conformations shown in (B). Note that the PC analysis involved using the 300-ps time-period not the whole 500-ns MD run. Inspired by the green curve, non-linear interpolation between Conf indexes #50 and #53 was used for conformational search and shown as black dots.

#### Essential motions of αB-cristallin57-69 revealed in the latent space

We selected 2 conformations in the latent space, Conf indexes #50 and #53, from **Figures 3A to C** to examine the conformational transitions in the latent space. Note that the MD saved one frame every 1 ps, and we re-saved a frame every 100 ps when preparing our training and validation sets. Therefore, Conf indexes #50 and #53 (2 red dots in **Figure 3D**) are significantly different; they present 2 endpoints from 300-ps time-period MD. To illustrate the conformation transition, we superimposed 4 conformations (indexes #50 to #53 **Figure 3A**) and found that the first 6 residues were maintained in a helix secondary structure, and the middle region around Leu9 showed minor but important fluctuations to twist the center of the peptide, whereas the flexible tail (residues 7 to 13) exhibited greater dynamics. A correlated sidechain movement (i.e., sidechains of Leu9 and Met12) was observed during the transitions as well.

In the latent space, we found that the 300-ps interval MD trajectory sampled conformational jiggling during the transitions (light blue dots in **Figure 3D**). To extract the essential motions from the conformation fluctuation, we applied PCA with BAT coordinates to analyze the major motions from the 300-ps time-period MD to quantify the motions. The first Principal Component (PC) revealed the major motions, which removed unessential fluctuations (**Figure 3B**). Presenting these essential motions in the latent space resulted in a very smooth curve (green dots in **Figure 3D**). Note that in the latent space, the PC motions appeared as a non-linear curve, not a straight line. **Figure S10C** exhibits smoother curves for motions following the second PC and the first + second PC modes, so the latent space recorded similar information obtained from PCA. As for PCA, our ICoN learned the important DOF that led to conformational transition (i.e., various sets of concerted dihedral rotations), and the latent space retained the information from DL. Therefore, interpolation between 2 dots in the latent space can efficiently sample protein motions that are led by movements from critical DOFs. As illustrated in **Figure 3**, the interpolation (black dots in **Figure 3D**) revealed the learned motions, which also showed a more rigid helix secondary structure in the first 6 residues and the fluctuations in the middle region around Leu9 to result in more distinct conformations with smooth transitions (**Figure 3C**). Importantly, our model accurately identified the concerted motions shown in both MD and PCA. These essential motions were in good agreement with those detected by dihedral PCA and our observation of the MD trajectory. Of note, our vBAT and model can precisely learn and present very small conformational changes, such as the methyl rotation of Met12.

#### Interpolation-generated synthetic conformations of αB-cristallin57-69

Because ICoN identified DOF that determines conformational changes with correct physics, we used the latent space interpolation as a conformational search engine to find more synthetic conformations. The search was achieved by interpolation between 2 dots with consecutive indexes (i.e., Conf indexes #10 and #11). Following the essential motions, we anticipated finding both new conformations and existing ones sampled by MD. Using the same non-linear interpolation function illustrated in **Figure 3D**, 10 points were generated for each pair, yielding 99,990 synthetic conformations from interpolating a total of 9,999 pairs of dots. We first eliminated high-energy conformations, then each new synthetic conformation was compared with its predecessors and eliminated if it was a repeat, yielding 43,484 distinct conformations (**Table 1**). We further compared these synthetic conformations with the raw MD data with 500,000 frames to identify 38,425 novel synthetic conformations.

We also examined the stability of the synthetic conformations quantified by their conformational energies using the molecular mechanics/generalized Born and surface areas (MM/GBSA) calculations. Conformations sampled from MD and our deep learning model have similar energy distribution (**Figure 4A**), which validates our synthetic conformations as being thermodynamically stable. In addition, we also compared our synthetic conformations with conformations obtained by another MD run (**Table 1**, Seed 2); none of the conformations from this MD was used in our training set. Our interpolation sampling found similar conformations found by another MD run **Figure 4B**, which demonstrates the efficiency and accuracy of the use of the latent space for conformational sampling.

**Figure 4.**
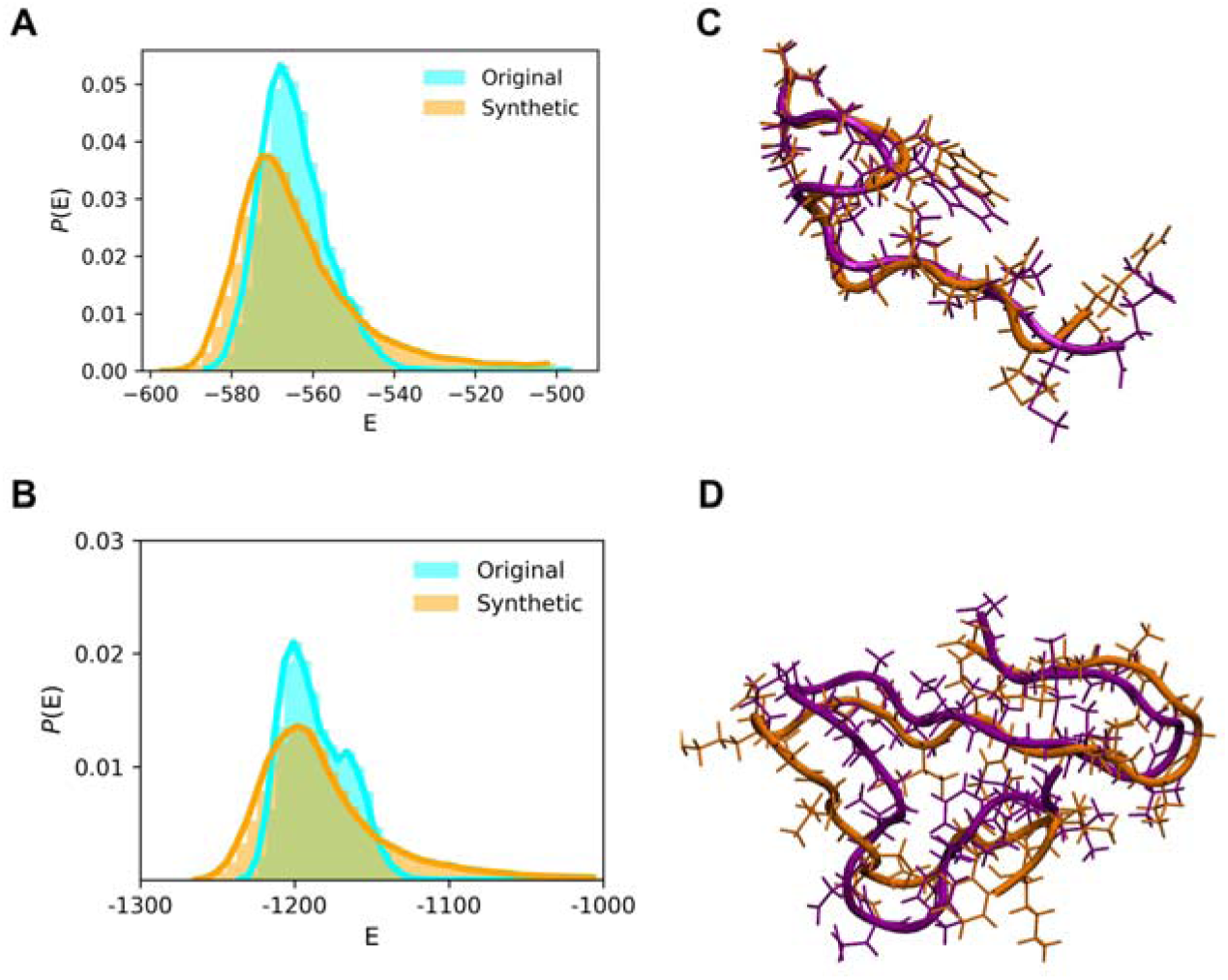
Examination of the stability of synthetic conformations using energy distribution and known conformations. Comparison of energy distribution from conformations of the original MD run (cyan) and synthetic (orange) conformations for (A) αB-crystallin57-69 and (B) Aβ42. Synthetic conformation (orange) similar to a known conformation sampled by other MD runs (purple) for (C) αB-crystallin57-69 and (D) Aβ42. The MD runs used for comparison but not used for training are Seed 2 and MD Run1.1-3 in Table 1.

### Latent space interpolation-generated synthetic conformations of Aβ42

The Aβ42 monomer exhibits a broad spectrum of conformations, from random coil to more structured α-helical and β-sheet conformations. Sampling strategies that can accurately model sidechain motions are crucial because the sidechain arrangements govern the intramolecular attractions that lead to various local turns and pre-organized shapes for subsequent oligomerization. Using the same hyperparameters obtained from MD Run1, we used the same strategy for αB-cristallin57-69 to interpolate 2 consecutive dots in the latent space for MD Runs 1 to 5 (**Table 1**) to obtain synthetic conformations. The 5 MD runs were initiated by a different Aβ42 monomer structure and/or with a different force field. To elucidate the search efficacy, we plotted the distribution with coordinates of Rg and RMSD (**Figure 5**) for MD Run1-5 and our synthetic conformations from Model 1-5 using these MD runs for training. The search using generative sampling in ICoN efficiently found many new conformations without the need for lengthy MD simulations (**Table 1)**.

**Figure 5.**
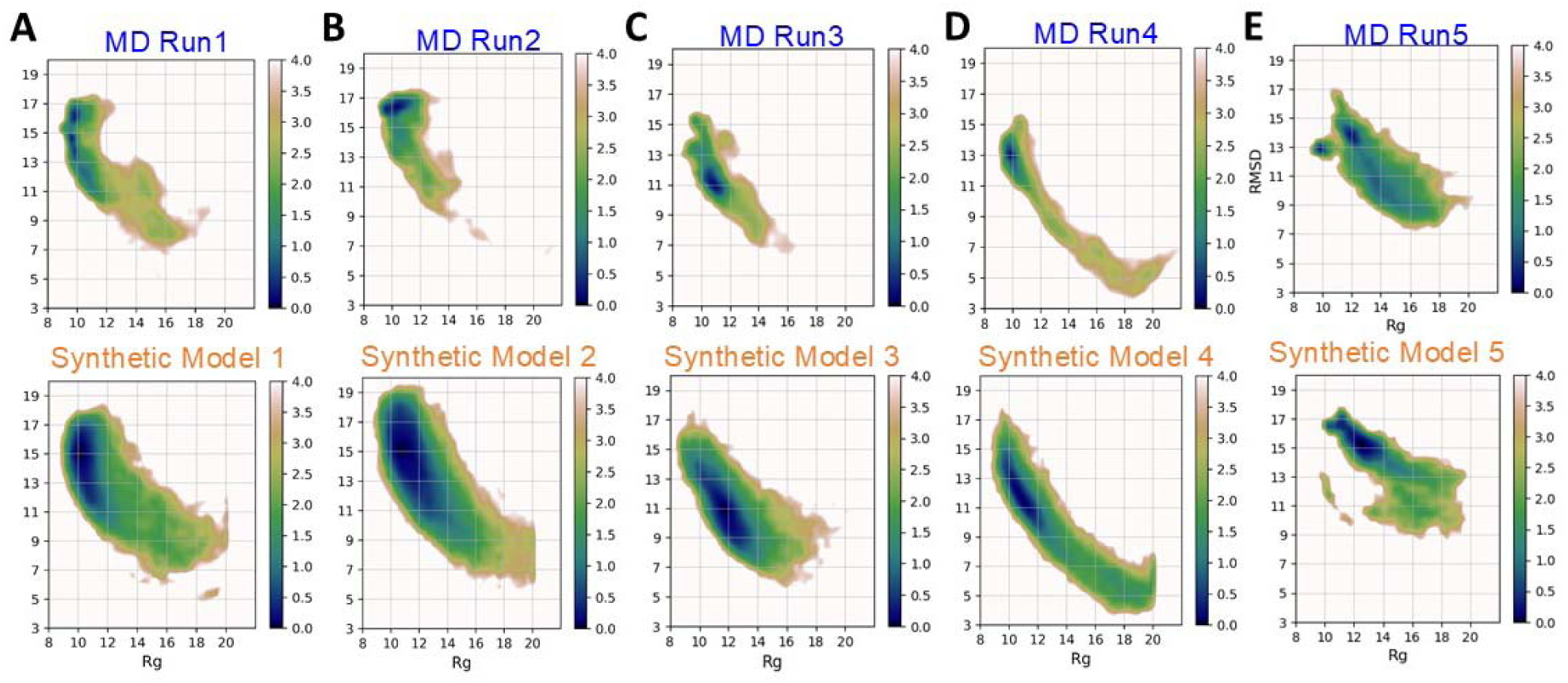
Free energy profile on the 2D space of RMSD (y-axis) and radius of gyration (Rg, x- axis) for Aβ42. For each pair of comparisons, the top depicts the conformation distribution from the MD run with frames are resaved every 10 ps and the bottom shows the distribution from distinct synthetic conformations. Using the same hyperparameters obtained from dataset MD Run1, Synthetic Models 1-5 was trained with MD frames obtained from MD Run1-5 (A to E), respectively. MD Runs 1 and 2 used the ff14IDPSFF, MD Runs 3 and 4 used the classical Amber ff14SB, and MD Run5 used EFFS1 force field. Notably, a highly packed initial conformation was used in MD Run 5 (**Figure S11**). The units of the free-energy landscapes are kcal/mol.

To validate that our synthetic conformations are thermodynamically stable, we again plotted the energy distribution of synthetic conformations found in Model 1 (**Figure 4B**). We also compared the synthetic conformations to another 3 MD runs using the same initial structure and IDP force field (MD Run1.1-1.3) to examine whether our search could find those sampled by other MD simulations. Although the Aβ42 monomer has numerous conformations, we could identify similar conformations (**Figure 4C**). In addition to checking the total conformational energy, we compared each energy component to verify that both bonded and non-bonded terms have energy distributions similar to those modeled by MD runs (**Figures S12 and S13**). Notably, the energy distribution from the synthetic conformations is relatively broader, thereby suggesting a greater diversity in the sampled conformations, which cover high and low-energy regions. The energy comparison underscores the high quality of structural integrity of the synthetic conformations.

### Analysis and biological implications of novel synthetic Aβ42 conformation ensembles

Obtaining Aβ42 monomer conformations is a critical step for understanding the mechanisms of initial encounters and interactions between monomers, which lead to subsequent oligomerization and fibrillization. Because the oligomerization steps can be highly sensitive to different environments (e.g., different membranes, existing fibrils, or ion concentrations), substantially different monomer conformations may initiate aggregation using different mechanisms in various environments. Several experiments and modeling work also suggest that some conformations are prone to be aggregated or non-toxic ^49–53^. Because of various experimental results regarding the monomer structure-function relationship in Aβ42 aggregation, we focus on structures with salt bridges and R5-A42 contacts to demonstrate the utility of the conformational search results from our generative AI model.

Of note, because our method efficiently sampled numerous low-energy monomer conformations, we used synthetic conformations found using Model 1 (**Table 1**) to demonstrate the biological importance of these conformations revealed by our ICoN model.

#### Major conformation turns into a global tertiary structural ensemble

The 127,673 distinct conformations (**Table 1**) are presented in the pairwise residue contact map (**Figure 6A**). Four local bends are marked as turn A (F4-H6), turn B (E11-H14), turn C (S26- K28), and turn D (V36-G38). We found novel synthetic conformations with all 4 turns (**Figure 1D**). Turn C has been widely reported in the fibril structures of both Aβ40^56^ and Aβ42^57^, and is the turn in the commonly referenced “β-turn-β” motif. Cryo-EM studies of brain-derived Aβ42 fibrils^57^ reveal that Aβ42 adopts an S-shaped fold with turns C and D being the two bends in the letter “S”. Turns A and B are located near the N-terminal, which is highly flexible and believed to be the metal binding region under abnormal physiological conditions (residues D1 to K16) ^58^. Although the conformations/turns and their population are obtained using only Model 1, they provide an overview of spatial arrangements of the Aβ42 monomer. Notably, although Aβ42 is intrinsically disordered, the protein sequence still can lead to many low-energy and highly populated conformations. Turn A located in the highly flexible N-terminus, where mutation substitutions of Arg5 suppressed the aggregation of Aβ42 ^50^. Our EPR measurements also showed distinct spectral features at residue 5, indicating that Arg5 may feature a distinct structure or interactions as compared with the nearby residues (**Figure 7**). The novel synthetic conformations revealed that Arg5 can form interactions with Ala42 with or without the presence of Turn A, as illustrated in **Figures 6B** and **6C**, respectively. Of note, turn D (V36-G38) is present alongside, which may help to orient and stabilize the C-terminus to form an interaction with Arg5 (red circle in **Figures 6B-C**). Our previous work showed that substituting a nitroxide spin label compound R1 at positions G37 and G38 altered the kinetics of Aβ42 fibril formation ^59^, and these synthetic conformations provide a possible monomer structure to guide future experimental design.

**Figure 6.**
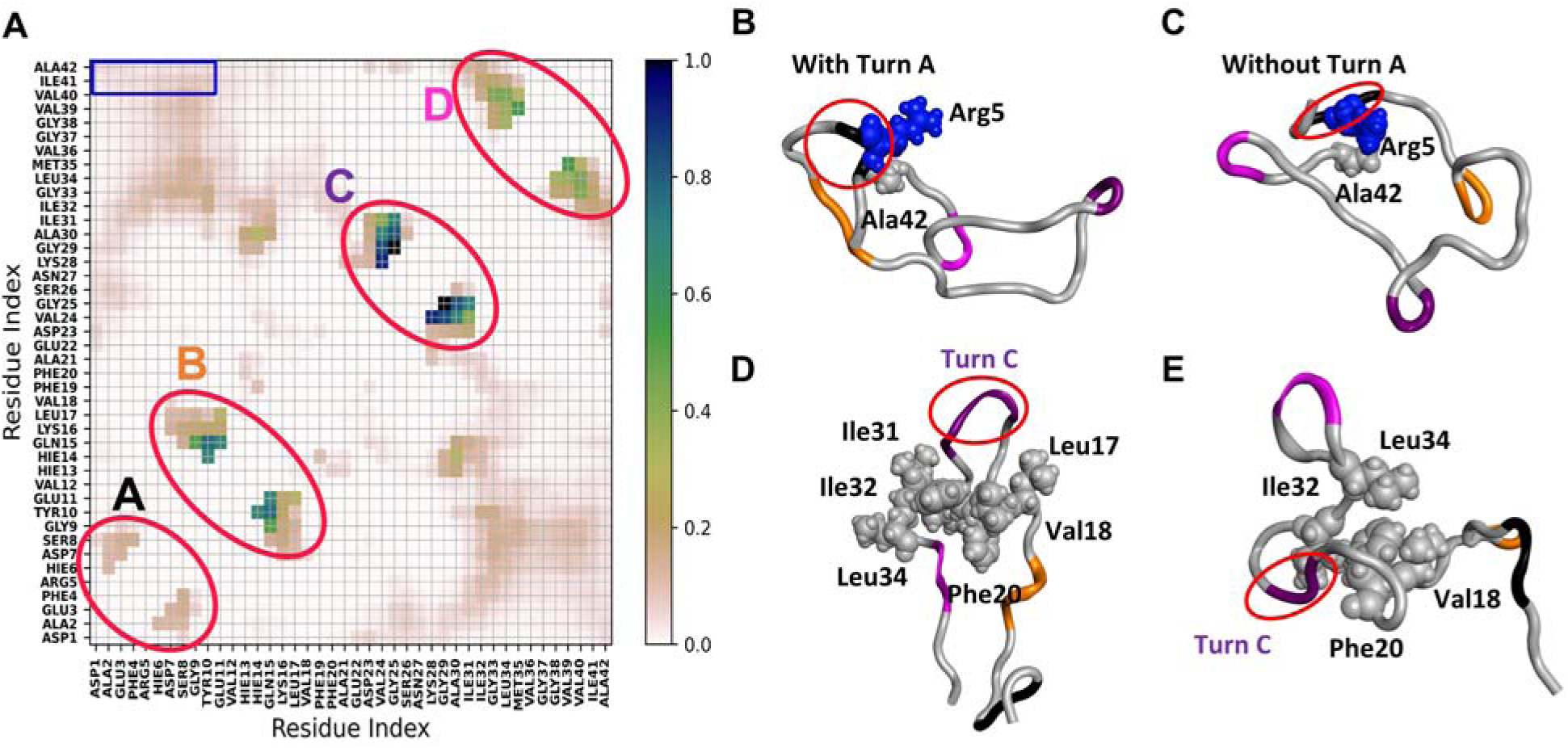
Probability of intramolecular contacts of synthetic conformations between Cα atoms of Aβ42 and representative conformations with Arg5-Ala42 contacts. **(A)** Four local bends are circled: turn A (F4-H6), turn B (E11-H14), turn C (S26-K28), and turn D (V36-G38). Residues contacting the C-terminal residues are marked in blue squares. Two representative novel synthetic conformations found in ICoN Model 1 with IDP force field depicting non-polar contacts between Arg5-Ala42 **(B)** with turn A and **(C)** without turn A. Both conformations utilize turn D to stabilize the C-terminus. **(D)** A representative novel synthetic conformation found in Model 3 with FF14SB force field shows an intermediate structure of forming Aβ42 tetramer which comprises a six stranded β-sheet with Turn C and hydrophobic core (L17-F20 and I31-L34). **(E)** A representative novel synthetic conformation from Model 5 with ESFF1 force field shows a structure with Turn B, C and D contributing to the formation of hydrophobic core. Residues of selection are shown in the van der Waal representation. Positively charged Arg and non-polar residues are blue and grey, respectively.

**Figure 7.**
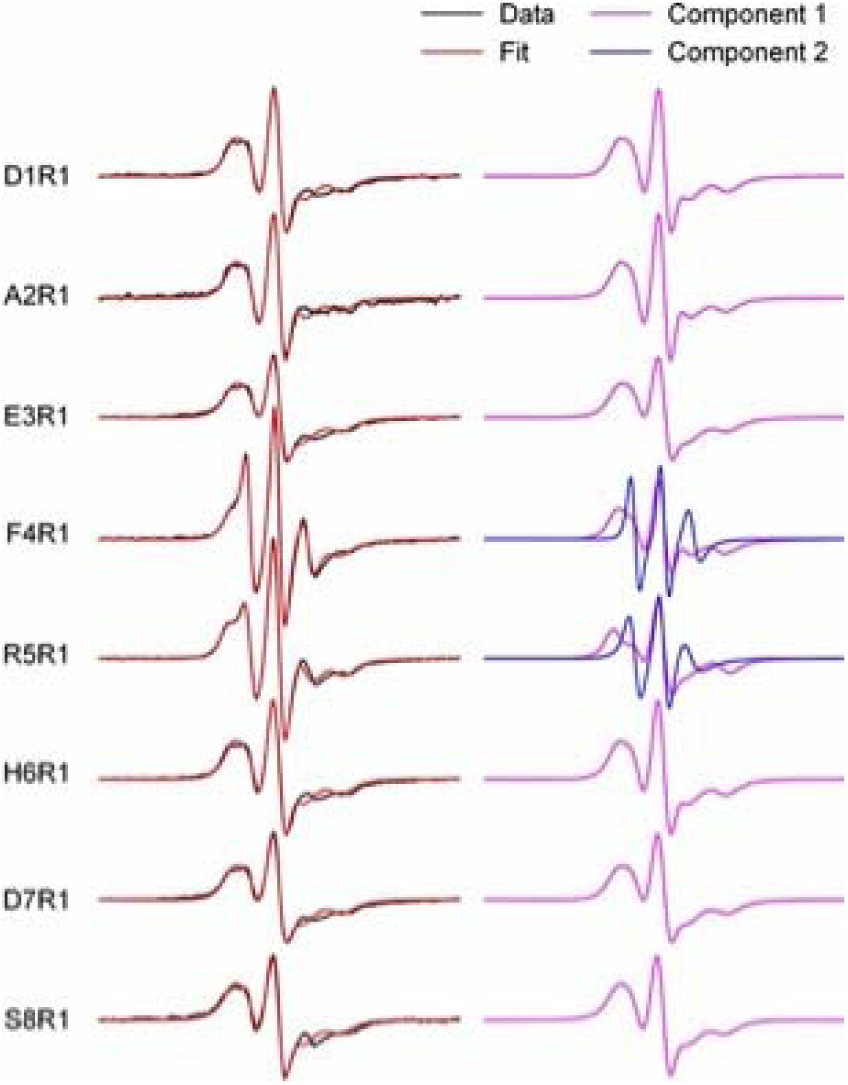
EPR spectra of spin-labeled Aβ42 fibrils show unique spectral features at residues 4 and 5. R1 represents the spin label. Left: EPR data are superimposed on the best fit of simulated EPR spectra. Right: Individual EPR spectral components for obtaining the best fit. Note that only the EPR spectra at residues 4 and 5 require a second spectral component. These data are consistent with a turn structure at residues 4-5.

Existing studies showed that Aβ42 monomer has numerous conformations, and MD simulations using initial conformation and/or force field can sample different conformations with few repeats ^42,43,47,60^. Using the hyperparameters obtained from **MD Run 1** to train datasets with conformations different from **MD Run2-5**, new intermediate states important in oligomerization were identified with ICoN. For example, as illustrated in **Figures 6 and 8**, novel synthetic conformations from the FF14SB force field (**Model 3 in Table 1**) reveal an intermediate state with a partial hairpin conformation which only requires minimal conformational arrangement to form and experimentally determined Aβ42 fibril from human brain ^57^ (**Figure 8A**). The partial hairpin conformations pre-organize hydrophobic regions ranging from L17-F20 and I31-L34 with residues F19 or F20 rotate inward to stabilize a hydrophobic core with surrounding nonpolar residues (**Figure 8B-C**). Interestingly, different directions of F19 and F20 could contribute to different types of brain-isolated fibril structures ^57^, and our ICoN model suggested the critical roles of their rearrangements. In addition, another partial hairpin conformation that has the potential of forming an Aβ42 tetramer structure, which comprises a six-stranded β-sheet with 2 β-strands in the middle and 2 β-turn-β on the side ^61^, is also observed (**Figure S14**). In Aβ42 tetramer, sidechains of F19 and F20 need to rotate outward to interact with surrounding monomers (**Figure S14A**). The partial hairpin conformation at Turn C leads to the formation of fibril conformations (**Figure 6D, 8, and S14**). Although the models were constructed using MD simulations for monomer, our synthetic conformations suggest a highly plausible intermediate state during fibril formation. Furthermore, novel synthetic conformations from ESFF1 force field (**Model 5 in Table 1**) show that turn C and turn D create highly packed hydrophobic regions prone to oligomerization. Specifically, we observed that V18, F20, I32, and L34 form a hydrophobic core (**Figure 6E**). Notably, unlike hydrogen bonds which are highly specific with additional geometry restraints, hydrophobic regions usually allow fluid-like sidechain movements to retain their flexibility. This different partial hairpin conformation from **Figure 6D** again preorganized the hydrophobic regions and preserved conformational plasticity for oligomerization.

**Figure 8.**
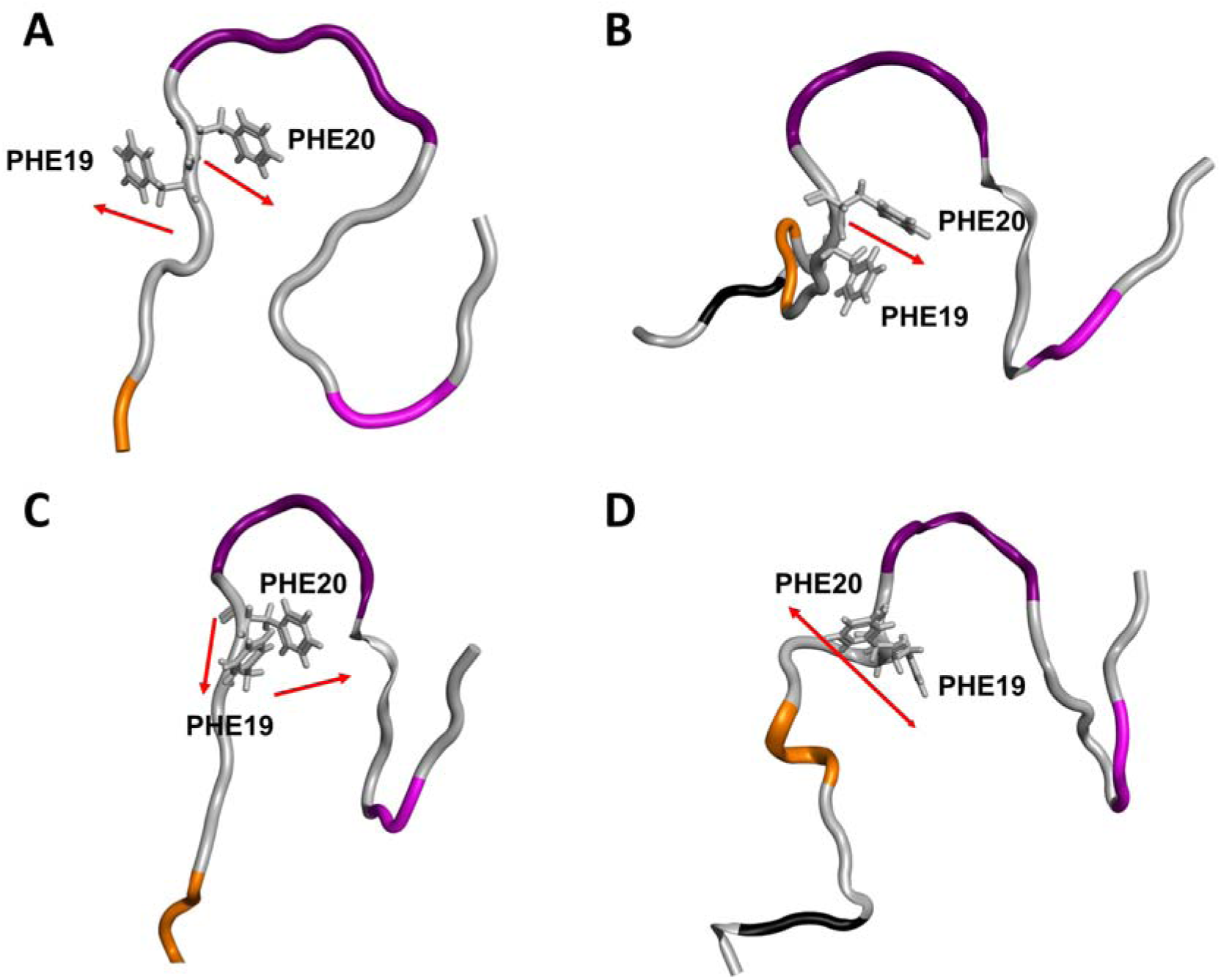
Intermediate state with partial hair pin at Turn C conformations which require minimal conformational arrangement to form Aβ42 fibril. Different orientations (direction indicated with red arrow) of F19 and F20 are observed leading to the formation of fibril conformation. **(A)** Crystal structure of hair pin conformation that form Aβ42 fibril from human brain (PDB: 7Q4M). **(B-D)** Novel sytnthetic strucutres are obtained from Model 3 with FF14SB force field. (**B**) Type I Aβ42 fibril with F19 and F20 rotate inward to form hydrophobic pocket. **(C)** Type II Aβ42 fibril with F20 rotate inward and F19 rotate outward. **(D)** Intermedicate states with F19 and F20 both rotate outside of the hydrophobic pocket.

#### Presence of salt bridges in toxic and less toxic monomer conformations

Salt bridges play an important role in providing intra-molecular attractions and conformational specificity that directly relate to Aβ42 aggregation. This non-covalent interaction forms when sidechains of oppositely charged residues are close enough to each other to experience electrostatic attractions. Therefore, protein structures must be described with precise atomistic details and sidechain motions sampled accurately. Although the D23-K28 salt bridge did not exist in the 1-µs MD Run1 used in our training set, by using interpolation in the latent space, the model sampled novel synthetic conformations with a new salt bridge, D23-K28 (**Figure 9**), which is reported to promote aggregation ^49^. Experiments suggested the importance of this salt bridge, but the conformation ensemble was never determined experimentally. The novel synthetic conformations revealed distinctly different rearrangements with the D23-K28 salt bridge, ranging from more extended to highly packed conformations (**Figure 9**). The D23-K28 constraint results in Turn C, which is important for oligomerization.

**Figure 9.**
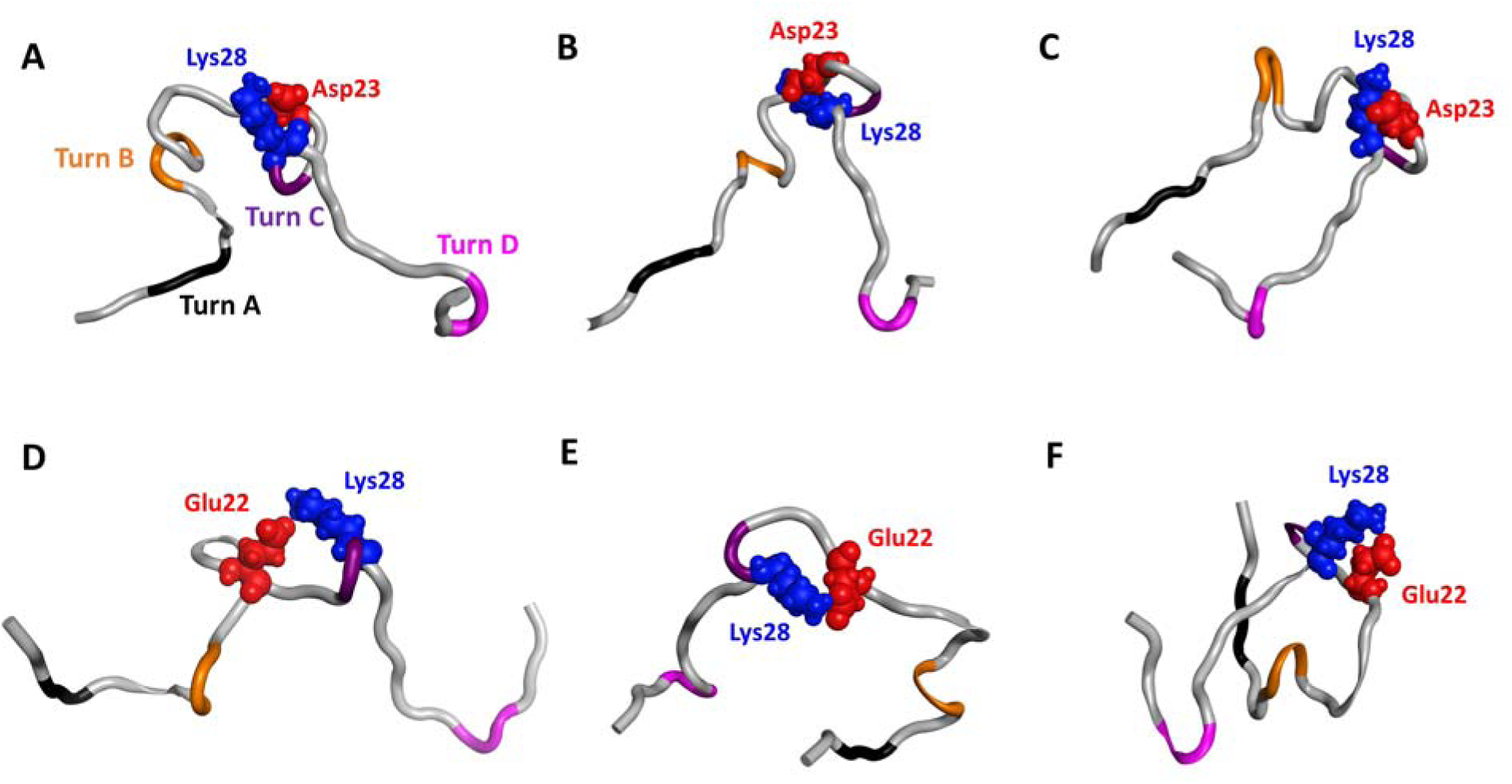
Novel synthetic conformations with intramolecular salt bridges. Top: structures with D23-K28 salt bridge. (A) and (B) show an open form with different local arrangements near turns B and C, and (C) reveals a U-shape, partial hairpin conformation. Bottom: structures with E22-K28 salt bridge with open (D), partial open (B) and compact (C) conformations.

The synthetic conformations revealed novel structures with the salt bridge E22-K28 (**Figures 9D-F**). This salt bridge was reported to prevent a toxic turn at positions E22 and D23, for potentially a less-toxic monomer ^62^. The synthetic conformation presents key features suggested by experiments, with no turn at positions 22 and 23 when the salt bridge E22-K28 exists (**Figure 9**). Nevertheless, other publications suggested that the salt bridge E22-K28 may help form other intramolecular contacts (e.g., contacts between the C-terminus and Arg5) to stabilize a hairpin structure and promote dimer formation ^50^. Although which salt bridges promote or inhibit monomer aggregation is debatable, our novel synthetic conformations provide various possible sidechain rearrangements and salt-bridge conformations for further interpreting experiments in understanding aggregation mechanisms.

#### Comparing ICoN with Phanto-IDP

Here we compare the conformational sampling using ICoN and Phanto-IDP, a new DL model developed by Chen’s^47^ which can efficiently sample backbone conformations of IDPs. Using backbone-only representation, Phanto-IDP utilizes a graph-based encoder and a variational layer to describe protein features. The variational layer sample neighboring area of points in latent space with mean, standard deviation, random variable and temperature to yield impressive sampling results with high diversity in a recent publication^47^. In this paper, the Chen group used MD Run 5 to train and generate new conformations with Phanto-IDP, where the raw data had backbone-only representation. For comparison, we added sidechains to their synthetic backbone conformations and carried out minimization and subsequent steps to obtain distinct newly synthetic conformations. Using the same MD Run 5, we applied the hyperparameters obtained from our MD Run 1 to sample new synthetic conformations with ICoN (Table 1). **Figure S15** presents a similar plot as **Figure 7** to compare the free energy profile on the RMSD and Rg for Aβ42 for conformations from MD and the novel synthetic ones. Using the graph-based representation for Aβ42 backbone, Phanto-IDP demonstrated great reconstruction accuracy (**Table S1**) and conformational sampling. Due to the nature of the use of mean and standard deviation in the variational layer, Phanto-IDP focuses on searching for new conformations close to conformations that are highly populated in an MD simulation. In contrast, ICoN samples conformations by interpolating data points in the latent space without discriminating popular conformations or rare events modeled by MD, resulting in a broad range of novel synthetic conformations. As illustrated in **Figure S16**, ICoN identified many new conformations in regions shown as transient states in MD Run5. These newly sampled sidechain rearrangements form a hydrophobic core with Turns B, C, and D, where similar hydrophobic regions and turns have been observed in several neurotoxic conformations determined by experiments (**Figure 6E and S16**). These results display the strength that using internal coordinates to represent both protein backbone and sidechains allows efficient combination of dihedral rotations in the latent space to sample a wide variety of new and biologically relevant synthetic conformations.

## Discussion

In this work, we demonstrated the capability of a deep learning model to directly generate new protein conformational ensembles by learning the physics of protein motions. Our approach utilized a neural network trained with protein conformations from classical MD simulations with all-atom representation in explicit solvent. We saved frames from MD trajectories without further processing, reducing protein atomic coordinates to a 3D latent space, where each conformation is a data point that can be accurately reconstructed back into a protein structure.

ICoN uses nonlinear interpolation between 2 data points in the 3D latent space to guide the search for conformational changes, identifying natural molecular motions. The vector-based BAT (vBAT) feature overcomes problems with Cartesian coordinates that fail to smoothly present dihedral rotation during conformation sampling, which is crucial in sampling large-scale motions, especially in IDPs. vBAT also effectively handles dihedral periodicity. The model is highly efficient for identifying novel synthetic Aβ42 conformation ensembles not seen in the MD trajectory. These ensembles have different large-scale motions (i.e., new compact conformations) and detailed sidechain rearrangements such as forming a new salt bridge between D23 and K28.

The synthetic Aβ42 conformations sampled by the ICoN model provide a comprehensive view of the conformational landscape for Aβ42 monomers. For example, the ICoN model finds salt bridges involving residues E22 or D23 **(Figure 8)**, both of which are hotspots for familial mutations in Alzheimer’s disease. The ICoN model also identifies four local bends at residue positions 4-6, 11-14, 26-28, and 36-38 (**Figures 1D and 6**). Although all 4 turns have been previously observed in isolation, it is worth noting that the experimentally determined Aβ42 structures do not contain all these turns in the same structure. For example, turn D has been found in the brain-derived Aβ42 fibril structures^57^, but is absent in the Aβ42 oligomers formed in the presence of detergents^61^. The co-occurrence of different turns in the synthetic Aβ42 conformations may help decode the Aβ aggregation pathways that lead to the formation of specific oligomers and fibrils.

Training of the ICoN model requires sufficient protein conformations, so the DL can reveal natural motions of the protein to guide the search toward new low-energy structures. Since experimentally determined structures typically cover only a small proportion of the overall conformation ensembles, we rely on MD simulations. Force-field parameters used in MD may affect the performance in sampling. For example, the model trained by MD runs using a less flexible ff14SB yielded fewer novel synthetic conformations (**Table 1**). However, our training set did not need to cover the complete conformational distribution. As long as conformation fluctuation governed by key dihedral rotations can be included in the training set, the model can identify these key elements for guiding the conformational search to generate novel synthetic conformations.

Conceptually, ICoN performs as a conformational search method rather than a deterministic simulation method. Therefore, the search results do not directly reflect the equilibrium distributions of protein conformations. While structured proteins have highly stable global energy minima, IDPs exhibit numerous fluctuating heterogeneous conformations with similar energy. The IDPs can be highly sensitive to their environment, making it impractical to precisely reproduce physiological conditions in experiments or simulations. Therefore, thoroughly finding IDP conformational ensembles is more practical and useful instead of aiming for the population distribution in equilibrium under specific physiological environment.

Importantly, unlike structured proteins whose native structures are functionally important, meta-stable conformations of IDPs can be crucial in performing biological functions. For example, experiments showed that Aβ42 monomer aggregation has a rate-limiting nucleation-dependent polymerization process ^53^, which suggests that the highly populated monomer conformations might not be the ones that drive nucleation, and the monomers are not pre-organized as a ready-to-aggregate conformation. Therefore, as compared with the conformational search for structure proteins, thoroughly sampling meta-stable conformations for IDPs is critical. As demonstrated in this study, the novel synthetic Aβ42 monomer conformations provide atomistic details to support several experimental observations, which also bring insights into mechanisms of oligomerization.

Conformational changes of biomolecules follow principles of physics, which are consequences of different arrangements of atoms rotating around a set of single bonds, the internal degrees of freedom having the highest flexibility. Both PCA and NMA describe the natural motions of a molecule, which are commonly used to determine essential protein dynamics and guide conformational sampling. The two techniques are linear transforms that extract the most important elements in a data matrix, a covariance or Hessian matrix. However, real protein systems can have very complicated and higher-order correlations that have intrinsic nonlinear effects and may not be well described using standard PCA or NMA. Moreover, although PC space is commonly used for data analysis, there is no function to convert manipulated data points in PC space back to its atomistic protein structure. The activation function used in the neural network adds nonlinear effects to process nonlinear features. With the use of BAT, a transition between two proteins can be achieved by interpolating two data points (conformations) in the latent space. As illustrated in **Figure 4**, if one obtains conformations along the smooth green curve in the latent space, the conformational changes reproduce the motions following the first PC mode. As for the natural motions identified by PCA or NMA, the deep learning algorithm learns the natural motions of the protein system (black dot line in **Figure 4**), and new conformations can be found by processing the interpolations in the latent space.

Although the ICoN model is fast, taking a few minutes to train 10,000 frames of Aβ42 and hours to perform quick energy minimization for the synthetic conformations generated from latent space interpolation using a GPU card, the training set is from a physics-based MD sampling. It typically takes days, if not weeks, to perform sufficiently long MD simulations to obtain the training set. To speed up the sampling, one can apply the ICoN model for preliminary results to select dissimilar protein conformations to seed more MD runs ^19^. An advantage in using BAT-based coordinates is that one can easily select a set of dihedral angles that determine the motions of interest for training, instead of using all DOF. For small proteins or peptides such as Aβ42 and αB-crystallin57-69, our classical internal BAT (Z-matrix)-based vBAT nicely presents local sidechain rearrangements and large-scale backbone motions. However, for larger (>200 residues) proteins, rotations of some dihedral angles near the root atoms may result in unrealistic motions on the far end of the protein. The issue can be addressed easily by using multilayer BAT coordinates, assigning multiple fragments/chains for a protein or multi-protein complex. Also, reconnecting fragments with pseudo-DOF can avoid the accumulated dependence problem ^63^. Future work will implement the multilayer BAT coordinates to eliminate the protein size limitation when building the ICoN model. Various interpolation and extrapolation strategies in the latent space will be examined for generating novel synthetic conformations as well.

## Methods

### All-atom Molecular Dynamics Simulations and Preparation of Training and Validation Data Sets

MD run1-4 were performed using the AMBER20 package using either ff14SB or ff14IDPSFF force fields (Table 1) ^64,65^. The systems were simulated using TIP3P explicit solvent model ^66^ at temperature of 298K with NPT ensemble. 12 Å cutoff was used for short range non-bonded interactions and the long-range electrostatic interactions were computed by the particle mesh Ewald method (PME) ^67^. MD run5 with ESFF1 force field, another IDP specific force field, is obtained from Chen’s group ^47,68^.

#### Aβ42

MD Run1 was initiated using PDB: 2NAO with ff14IDPSFF force field (**Table 1**). Each frame was saved at 1-ps time integral which made up a total of 1,000,000 frames. Other MD trajectories using either PDB 2NAO or 1Z0Q as the initial structure were obtained from our previous work and detailed in Table 1 ^54,55,69^. Conformations of **MD Run1 to 5** were used for training, validation, sampling conformations from latent space and generate synthetic conformations, while **MD Run1.1 to 1.3** were utilized for pairwise RMSD search for synthetic conformations obtained by Model 1 (**Table 1**).

#### aβ-crystallin57-69

AlphaFold2 computed structure of aβ-crystallin57-69 (PDB: AF_AFP02511F1) was used as initial structure for MD simulations. Two randomly seeded 500-ns MD simulations were obtained from our previous work, where each frame was collected at 1-ps interval which made up 500,000 frames ^70^. Conformations of Seed1 (**Table 1**) were used for training, validation, and sampling conformations from latent space, while Seed2 was utilized for pairwise RMSD search for synthetic conformations.

#### Preparation of training and validation data sets

To construct our dataset for ICoN model training, we saved 1 frame every 100-ps starting from 0-ps of Aβ42 MD Run1-5 (10,000 conformations; 1% of raw MD) to prepare training sets.

Similarly, we saved 1 frame every 100-ps starting from frame 50-ps of Aβ42 MD Run1-5 for validation sets. Similar approach is used for aβ-crystallin57-69 Seed1, where 1 frame every 50- ps starting from 0-ps (5,000 conformations; 1% of raw MD) used for training, and 1 frame every 50-ps starting from 25-ps used for validation. This ensures training and validation datasets are separated by 50-ps and 25-ps time interval for Aβ42 and aβ-crystallin57-69 respectively. Details of other training and validation datasets are shown in Table 1. Due to the highly dynamic nature of Aβ42, every 100ps of the MD simulations will yield distinct conformations, and thus using the 10,000 conformations are enough to capture the physics. This approach allows training a model from minimal amount of input data yet generate numerous conformations with extended conformational space.

#### Dihedral PCA (dPCA)

To extract conformations with major protein motions, we utilized dihedral Principal Component Analysis (dPCA) employing torsion angles with in-house code. The resulting conformations, derived from the first three principal components, were then projected onto a latent space using the IcoN model in order to get insight on conformational transition of the aβ-crystallin57-69 300- ps trajectory, composed of 300 frames. Notably, the transitions observed in the latent space, governed by principal component modes, manifested as smooth nonlinear trajectories. Consequently, we opted for nonlinear interpolation to produce synthetic conformations.

The dPCA calculation involves mapping torsion angles to a unit circle using trigonometric functions. This enables accurate estimation of differences (involving subtraction) and averages (involving summation) during the construction of the covariance matrix, thereby preventing erroneous computation of their correlation at the discontinuity margin (±180° or 360°/0°) ^63,71,72^.

### Protein Structure Representation

#### Classical Bond-Angle-Torsion coordinates

Bond Angle Torsion (BAT) is the internal coordinate representation ^73–75^, where “bond” denotes the distance between a pair of bonded atoms, “angle” refers to the bond angle between pairs of bonds connected to a central atom, and “torsion” indicates the dihedral angle formed by four bonded atoms via three bonds. Specifically, the dihedral angle is defined as the angle between the plane containing atoms (i, j, k) and another plane containing atoms (j, k, l) (**Figure 1E**). BAT coordinates can be accurately transformed back to all atom Cartesian coordinates. In this, the placement of each atom i > 3 that is not bounded to atom 2 is specified by its bond length (b_i_), bond angle (α_i_), and dihedral angle (θ_i_) with respect to three other atoms that are bonded in sequence and whose positions are already defined (**details in SI**). Three terminal atoms, designated as root-based atoms, are used to initiate conversion from BAT to Cartesian. Root based atoms carry six external degrees of freedom such as global translations and rotations.

#### Vector Representation of BAT (vBAT)

Although BAT coordinates effectively capture concerted motions, they suffer from a periodicity problem. Dihedral rotation from 179° to-179° is actually a 2° shift in angular space. However, the basic arithmetic operation used in computing a covariance matrix results in a large, 179°-(- 179°) = 358°, rotation ^71^. This issue can be addressed by mapping dihedral angles into circular space using trigonometric functions, though this comes with additional computational cost.

However, this cost can be avoided by using vBAT as input/output features. Hence, we used vector representation of BAT coordinates termed vBAT as input features to the ICoN model. The vBAT internal coordinate representation is equivariant under global translations and rotations, and thus provided strong inductive bias for our ICoN model. Furthermore, compared to other structural representations such as Anchored Cartesian ^42^ or PCA based representations ^33^, internal coordinate representation is independent of reference structure. A detailed description of vBAT computation is provided in this section and in Supplementary Information.

For a set of four bonded atoms with atom indexes i, j, k, and l (**Figure 2B**), we defined three bond vectors v_ij_, v_jk_, and v_kl,_ that are the relative positions between the atoms. The unit vectors that are normal to the plane made by a pair of neighboring bond vectors defined vector features, denoted as 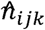 and 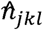.

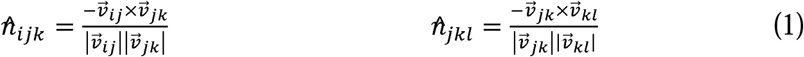

vBAT vector features comprising 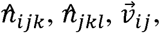 and 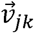 were used as an input to ICoN model.

When sampling conformations in latent space, all external degrees of freedom (root-based coordinates) were inferred from the reference conformation. Sampled features 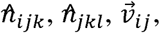 and 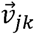 were first converted to bond angles, and torsions, and then transformed back to Cartesian coordinates in large batches using a highly optimized GPU implementation (**Figure S2**). Since changes in bond length are very minimal throughout MD simulations, we used fixed bond length inferring from the reference structure for all generated conformations.

### Model Architecture and Training

#### Model Architecture

The ICoN model is an autoencoder type architecture based on fully connected neural network (**Figure 2A**) trained on MD data to directly output all atom protein conformations. The model is designed to compress an input representation into lower dimensional latent space (3D) with an encoder that can be directly visualized. These points representing conformations in 3D latent space are decompressed back to the original input dimension with decoder.

For the first step, vBAT vectors were flattened followed by concatenation into a high dimensional array to use as input features to the ICoN model. As a result, we have 4×3×(N-3) features for each molecule 2376, and 7488 features for αB-crystallin57-69 and aβ42 respectively. The activation units were gradually reduced to 3-dimensional latent space in seven layers, where each molecular conformation can be presented in the 3D latent space, as illustrated in **Figure 2A**. For both systems, the encoding process reduced the dimension from n_f_, to n_f_/4, n_f_/8, n_f_/16, n_f_/32, n_f_/64, n_f_/64, and 3, where n_f_ is the total number of input features. Then, the decoding process brought the representation from 3D to 2376 dimension with the same number of layers but reverse the order.

The LeakyReLU activation function was applied to all layers except the last layer. Layer normalization is applied in the second layer. To prevent overfitting, 10% of the weights were randomly set to zero in the third layer, which led to better convergence of validation loss. The decoder takes the latent vector as input and decompresses it back to its original feature dimension. The decoder also featured LeakyReLU nonlinearity but did not include any layer normalization or dropout layers. The bias units are omitted aiming to minimize the model’s parameter count. Meanwhile, all model parameters were initialized through the Xavier Uniform weight initialization method, which is known to enhance training stability and efficiency ^76^. The total count of model parameters utilized is 3,294,964 for αB-crystallin57-69 and 32,721,156 for aβ42. Throughout the iterative training process, in addition to learning a relationship between vast number of features, the network also learns how to meaningfully compress the representation into the lower dimensional 3D latent space by performing nonlinear dimensionality reduction. Each conformation is converted from vBAT to Cartesian for further analysis.

#### Loss Function

The combination of L1 and L2 loss functions, the Smooth L1 loss, was used to minimize the difference between the model output (y_i_) and the original input (x_i_). The equation for the smooth L1 loss function, L, is given below (Eq. 2):

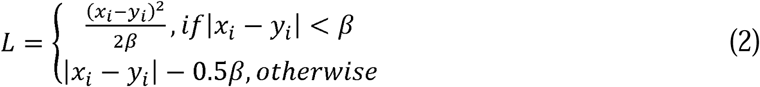

where, (x_i_-y_i_)^2^ is L2 loss and |x_i_-y_i_| is L1 loss.

The L1 and L2 functions intersect at β=1.0 resulting in a smooth, continuous loss function that is less sensitive to outliers compared to the L1 loss. However, for larger errors, the function transitions to an L1 loss, ensuring robustness to extreme values. As a result, using Smooth L1 loss can lead to more stable training and better convergence.

Optimization was performed using the Adam optimizer ^77^ for both protein systems where optimal learning rate found was 0.001 via hyper-parameter tuning (details in Figure S17). For Aβ42 training, the learning rate was manually reduced from 0.0002 to 0.0001 after 5000 iterations, resulting in a total of 10,000 training iterations. The learning rate was kept constant at 0.001 for aβ-crystallin57-69, comprising a total of 5000 training iterations. We fed 200 conformations in mini-batches per iteration to the model, randomly selecting frames from the data pool during each training iteration. Training was terminated upon convergence of both the training and validation loss, a process that took a few minutes for both systems. The MDAnalysis Python library ^78^ used as the tool of choice for reading and writing trajectories.

#### Evaluation of reconstructed conformations

Reconstruction accuracy was measured by computing the RMSD of all heavy atoms (all non-hydrogen atoms) and backbone atoms (N, Cα, C, and O atoms) between the original and reconstructed conformations (Table 1) (see SI for details).

#### Implementation

In this study, we employed the PyTorch library to implement all deep learning algorithms ^79^. MDAnalysis, a Python library, facilitated MD trajectory input/output operations and torsion list construction^78^. Leveraging the torch tensor data structure in PyTorch, we implemented GPU-optimized BAT and vBAT calculations. The code and pre-trained ICoN model parameters can be accessed at the following link: https://github.com/chang-group/ICoN

### Generation of Novel Conformations

In this process we have 4 steps. The first step is interpolation. We first encode training and validation data set into the latent space. Then, non-linear interpolation is performed between each pair of consecutive points in 3D latent space. 10 points were interpolated from each consecutive pair of points yielding a total of 19,9990 and 99,990 interpolated conformations for Aβ42 MD-Run1 and aβ-crystallin57-69-Seed1, respectively (**Table 1**). In the second step, we refine the generated structures through minimization to avoid potential atomic clashes. Steps three and four remove self-repeats and repeats with raw MD to ensure the creation of novel and unique synthetic conformations (see details of each step in SI).

### Analysis of synthetic conformations

#### Free energy profiles

To derive free energy profiles, we initially constructed a two-dimensional histogram by binning the radius of gyration (Rg) and backbone root mean squared deviation (RMSD) of protein conformations with respect to a reference structure. Subsequently, we determined the normalized population (P) for each bin throughout the entire trajectory. The free energy values were then computed using the equation F =-k_B_T log(P), where k_B_ represents the Boltzmann constant and T is a room temperature.

#### Contact maps

The contact maps were generated by applying a cutoff of 6.5 Å to all intramolecular distances between C-alpha positions. A contact density of one is assigned when a contact is formed, and zero otherwise, for each frame. Local contact densities of residue pairs with indexes i and j, where |i-j| < 4 were set to zero. The average across all frames (conformations) yielded the contact probability between the pairs of residues.

#### Aβ42 turn characterization

Four local turns have been identified as turn A (F4-H6), turn B (E11-H14), turn C (S26-K28), and turn D (V36-G38). To selectively isolate conformations characterized by turns occurring in specific regions, we initially identified residue pairs exhibiting a high pair contact density in proximity to the corresponding turn region, as depicted in contact map of **Figure 7**. Subsequently, a turn was deemed to be formed if the C-alpha distance between the selected pair of residues was less than 6.5 Å. Conversely, a turn was classified as not formed when the contact distance exceeded 9.0 Å. For turns A, B, C, and D, the residue pairs utilized for contact identification are Ala2:Asp7, Tyr10:Gly15, Val24:Gln29, and Gly33:Val40, respectively are selected.

## Data Availability

The code, pre-trained ICoN model parameters, and data sets are available at: https://github.com/chang-group/ICoN

## Acknowledgements.

We thank Dr. Hisashi Okumura and Dr. Mai Suan Li for providing their modeled Aβ42 conformations, Dr. Hai-Feng Chen and Junjie Zhu for providing their MD trajectory and generated structures of Aβ42 from Phanto-IDP, and Dr. David Minh and Ruben Montes for helpful discussion. This study was supported by the US National Institutes of Health (R01GM- 109045 to C. C. and R01AG-050687 and R21AG-082041 to Z.G.), the US National Science Foundation (MCB-1350401 to C.C.) and the Bourns Endowment funds to B.B.)

## Notes

### Competing Interest Statement

The authors have declared no competing interest.

### Summary of Updates

We corrected several grammatical error, clarified the sentences and included necessary citation and adjust the Figures.

## References

(1) Miller, M. D.; Phillips, G. N. Moving beyond Static Snapshots: Protein Dynamics and the Protein Data Bank. Journal of Biological Chemistry 2021, 296, 100749. 10.1016/j.jbc.2021.100749.

(2) Dill, K. A.; Bromberg, S. Molecular Driving Forces: Statistical Thermodynamics in Biology, Chemistry, Physics, and Nanoscience; Garland Science, 2011.

(3) Zhang, Y.; Chen, H.; Li, R.; Sterling, K.; Song, W. Amyloid β-Based Therapy for Alzheimer’s Disease: Challenges, Successes and Future. Signal Transduct Target Ther 2023, 8 (1), 248. 10.1038/s41392-023-01484-7.

(4) Huang, B.; Kong, L.; Wang, C.; Ju, F.; Zhang, Q.; Zhu, J.; Gong, T.; Zhang, H.; Yu, C.; Zheng, W.-M.; Bu, D. Protein Structure Prediction: Challenges, Advances, and the Shift of Research Paradigms. Genomics Proteomics Bioinformatics 2023, 21 (5), 913–925. 10.1016/j.gpb.2022.11.014.

(5) Whitford, D. Proteins: Structure and Function; Wiley, 2013.

(6) Tripathi, T.; Dubey, V. K. Advances in Protein Molecular and Structural Biology Methods; Elsevier Science, 2022.

(7) Macuglia, D.; Roux, B.; Ciccotti, G. The Emergence of Protein Dynamics Simulations: How Computational Statistical Mechanics Met Biochemistry. The European Physical Journal H 2022, 47 (1), 13. 10.1140/epjh/s13129-022-00043-y.

(8) Hollingsworth, S. A.; Dror, R. O. Molecular Dynamics Simulation for All. Neuron 2018, 99 (6), 1129–1143. 10.1016/j.neuron.2018.08.011.

(9) Allison, J. R. Computational Methods for Exploring Protein Conformations. Biochem Soc Trans 2020, 48 (4), 1707–1724. 10.1042/BST20200193.

(10) Paquet, E.; Viktor, H. L. Molecular Dynamics, Monte Carlo Simulations, and Langevin Dynamics: A Computational Review. Biomed Res Int 2015, 2015, 1–18. 10.1155/2015/183918.

(11) T. Daniel Crawford; Cecilia Clementi; Robert Harrison; Teresa Head-Gordon; Shantenu Jha; Anna Krylov; Ashley Ringer-McDonald; Theresa Windus; Dominika Zgid. Molssi. https://molssi.org/molssi-community-code-partners/ (accessed 2024-04-19).

(12) Bruce, N. J.; Ganotra, G. K.; Richter, S.; Wade, R. C. KBbox: A Toolbox of Computational Methods for Studying the Kinetics of Molecular Binding. J Chem Inf Model 2019, 59 (9), 3630–3634. 10.1021/acs.jcim.9b00485.

(13) David, C. C.; Jacobs, D. J. Principal Component Analysis: A Method for Determining the Essential Dynamics of Proteins; 2014; pp 193–226. 10.1007/978-1-62703-658-0_11.

(14) Yasuda, T.; Shigeta, Y.; Harada, R. Efficient Conformational Sampling of Collective Motions of Proteins with Principal Component Analysis-Based Parallel Cascade Selection Molecular Dynamics. J Chem Inf Model 2020, 60 (8), 4021–4029. 10.1021/acs.jcim.0c00580.

(15) Peng, C.; Wang, J.; Shi, Y.; Xu, Z.; Zhu, W. Increasing the Sampling Efficiency of Protein Conformational Change by Combining a Modified Replica Exchange Molecular Dynamics and Normal Mode Analysis. J Chem Theory Comput 2021, 17 (1), 13–28. 10.1021/acs.jctc.0c00592.

(16) Chang, C.; Gilson, M. K. Tork: Conformational Analysis Method for Molecules and Complexes. J Comput Chem 2003, 24 (16), 1987–1998. 10.1002/jcc.10325.

(17) Kolossváry, I.; Guida, W. C. Low Mode Search. An Efficient, Automated Computational Method for Conformational Analysis: Application to Cyclic and Acyclic Alkanes and Cyclic Peptides. J Am Chem Soc 1996, 118 (21), 5011–5019. 10.1021/ja952478m.

(18) Kanada, R.; Terayama, K.; Tokuhisa, A.; Matsumoto, S.; Okuno, Y. Enhanced Conformational Sampling with an Adaptive Coarse-Grained Elastic Network Model Using Short-Time All-Atom Molecular Dynamics. J Chem Theory Comput 2022, 18 (4), 2062–2074. 10.1021/acs.jctc.1c01074.

(19) Tian, H.; Jiang, X.; Xiao, S.; La Force, H.; Larson, E. C.; Tao, P. LAST: Latent Space-Assisted Adaptive Sampling for Protein Trajectories. J Chem Inf Model 2023, 63 (1), 67–75. 10.1021/acs.jcim.2c01213.

(20) Ghorbani, M.; Prasad, S.; Klauda, J. B.; Brooks, B. R. GraphVAMPNet, Using Graph Neural Networks and Variational Approach to Markov Processes for Dynamical Modeling of Biomolecules. J Chem Phys 2022, 156 (18). 10.1063/5.0085607.

(21) Lemke, T.; Peter, C. EncoderMap: Dimensionality Reduction and Generation of Molecule Conformations. J Chem Theory Comput 2019, 15 (2), 1209–1215. 10.1021/acs.jctc.8b00975.

(22) Merz, K. M.; Wei, G.-W.; Zhu, F. Editorial: Machine Learning in Bio-Cheminformatics. J Chem Inf Model 2024, 64 (7), 2125–2128. 10.1021/acs.jcim.4c00444.

(23) Bonati, L.; Zhang, Y.-Y.; Parrinello, M. Neural Networks-Based Variationally Enhanced Sampling. Proceedings of the National Academy of Sciences 2019, 116 (36), 17641–17647. 10.1073/pnas.1907975116.

(24) Mulnaes, D.; Gohlke, H. TopScore: Using Deep Neural Networks and Large Diverse Data Sets for Accurate Protein Model Quality Assessment. J Chem Theory Comput 2018, 14 (11), 6117–6126. 10.1021/acs.jctc.8b00690.

(25) Prašnikar, E.; Ljubič, M.; Perdih, A.; Borišek, J. Machine Learning Heralding a New Development Phase in Molecular Dynamics Simulations. Artif Intell Rev 2024, 57 (4), 102. 10.1007/s10462-024-10731-4.

(26) Kikutsuji, T.; Mori, Y.; Okazaki, K.; Mori, T.; Kim, K.; Matubayasi, N. Explaining Reaction Coordinates of Alanine Dipeptide Isomerization Obtained from Deep Neural Networks Using Explainable Artificial Intelligence (XAI). J Chem Phys 2022, 156 (15), 154108. 10.1063/5.0087310.

(27) Do, H. N.; Wang, J.; Bhattarai, A.; Miao, Y. GLOW: A Workflow Integrating Gaussian-Accelerated Molecular Dynamics and Deep Learning for Free Energy Profiling. J Chem Theory Comput 2022, 18 (3), 1423–1436. 10.1021/acs.jctc.1c01055.

(28) Zhu, J.; Li, Z.; Zhang, B.; Zheng, Z.; Zhong, B.; Bai, J.; Wang, T.; Wei, T.; Yang, J.; Chen, H.-F. Precise Generation of Conformational Ensembles for Intrinsically Disordered Proteins Using Fine-Tuned Diffusion Models. 10.1101/2024.05.05.592611.

(29) Majewski, M.; Pérez, A.; Thölke, P.; Doerr, S.; Charron, N. E.; Giorgino, T.; Husic, B. E.; Clementi, C.; Noé, F.; De Fabritiis, G. Machine Learning Coarse-Grained Potentials of Protein Thermodynamics. Nat Commun 2023, 14 (1), 5739. 10.1038/s41467-023-41343-1.

(30) Taneja, I.; Lasker, K. Machine-Learning-Based Methods to Generate Conformational Ensembles of Disordered Proteins. Biophys J 2024, 123 (1), 101–113. 10.1016/j.bpj.2023.12.001.

(31) Glielmo, A.; Husic, B. E.; Rodriguez, A.; Clementi, C.; Noé, F.; Laio, A. Unsupervised Learning Methods for Molecular Simulation Data. Chem Rev 2021, 121 (16), 9722–9758. 10.1021/acs.chemrev.0c01195.

(32) Zheng, L.-E.; Barethiya, S.; Nordquist, E.; Chen, J. Machine Learning Generation of Dynamic Protein Conformational Ensembles. Molecules 2023, 28 (10), 4047. 10.3390/molecules28104047.

(33) Noé, F.; Olsson, S.; Köhler, J.; Wu, H. Boltzmann Generators: Sampling Equilibrium States of Many-Body Systems with Deep Learning. Science (1979) 2019, 365 (6457). 10.1126/science.aaw1147.

(34) Ramaswamy, V. K.; Musson, S. C.; Willcocks, C. G.; Degiacomi, M. T. Deep Learning Protein Conformational Space with Convolutions and Latent Interpolations. Phys Rev X 2021, 11 (1), 011052. 10.1103/PhysRevX.11.011052.

(35) Atz, K.; Grisoni, F.; Schneider, G. Geometric Deep Learning on Molecular Representations. Nat Mach Intell 2021, 3 (12), 1023–1032. 10.1038/s42256-021-00418-8.

(36) Jin, Y.; Johannissen, L. O.; Hay, S. Predicting New Protein Conformations from Molecular Dynamics Simulation Conformational Landscapes and Machine Learning. *Proteins: Structure*, Function, and Bioinformatics 2021, 89 (8), 915–921. 10.1002/prot.26068.

(37) Audagnotto, M.; Czechtizky, W.; De Maria, L.; Käck, H.; Papoian, G.; Tornberg, L.; Tyrchan, C.; Ulander, J. Machine Learning/Molecular Dynamic Protein Structure Prediction Approach to Investigate the Protein Conformational Ensemble. Sci Rep 2022, 12 (1), 10018. 10.1038/s41598-022-13714-z.

(38) Sultan, M. M.; Wayment-Steele, H. K.; Pande, V. S. Transferable Neural Networks for Enhanced Sampling of Protein Dynamics. J Chem Theory Comput 2018, 14 (4), 1887–1894. 10.1021/acs.jctc.8b00025.

(39) Mehdi, S.; Smith, Z.; Herron, L.; Zou, Z.; Tiwary, P. Enhanced Sampling with Machine Learning. Annu Rev Phys Chem 2024, 75 (1). 10.1146/annurev-physchem-083122-125941.

(40) Hoseini, P.; Zhao, L.; Shehu, A. Generative Deep Learning for Macromolecular Structure and Dynamics. Curr Opin Struct Biol 2021, 67, 170–177. 10.1016/j.sbi.2020.11.012.

(41) Ward, M. D.; Zimmerman, M. I.; Meller, A.; Chung, M.; Swamidass, S. J.; Bowman, G. R. Deep Learning the Structural Determinants of Protein Biochemical Properties by Comparing Structural Ensembles with DiffNets. Nat Commun 2021, 12 (1), 3023. 10.1038/s41467-021-23246-1.

(42) Gupta, A.; Dey, S.; Hicks, A.; Zhou, H.-X. Artificial Intelligence Guided Conformational Mining of Intrinsically Disordered Proteins. Commun Biol 2022, 5 (1), 610. 10.1038/s42003-022-03562-y.

(43) Janson, G.; Valdes-Garcia, G.; Heo, L.; Feig, M. Direct Generation of Protein Conformational Ensembles via Machine Learning. Nat Commun 2023, 14 (1), 774. 10.1038/s41467-023-36443-x.

(44) Teixeira, J. M. C.; Liu, Z. H.; Namini, A.; Li, J.; Vernon, R. M.; Krzeminski, M.; Shamandy, A. A.; Zhang, O.; Haghighatlari, M.; Yu, L.; Head-Gordon, T.; Forman-Kay, J. D. IDPConformerGenerator: A Flexible Software Suite for Sampling the Conformational Space of Disordered Protein States. J Phys Chem A 2022, 126 (35), 5985–6003. 10.1021/acs.jpca.2c03726.

(45) Zhu, J.-J.; Zhang, N.-J.; Wei, T.; Chen, H.-F. Enhancing Conformational Sampling for Intrinsically Disordered and Ordered Proteins by Variational Autoencoder. Int J Mol Sci 2023, 24 (8), 6896. 10.3390/ijms24086896.

(46) Ramanathan, A.; Ma, H.; Parvatikar, A.; Chennubhotla, S. C. Artificial Intelligence Techniques for Integrative Structural Biology of Intrinsically Disordered Proteins. Curr Opin Struct Biol 2021, 66, 216–224. 10.1016/j.sbi.2020.12.001.

(47) Zhu, J.; Li, Z.; Tong, H.; Lu, Z.; Zhang, N.; Wei, T.; Chen, H.-F. Phanto-IDP: Compact Model for Precise Intrinsically Disordered Protein Backbone Generation and Enhanced Sampling. Brief Bioinform 2023, 25 (1). 10.1093/bib/bbad429.

(48) Toyama, B. H.; Hetzer, M. W. Protein Homeostasis: Live Long, Won’t Prosper. Nat Rev Mol Cell Biol 2013, 14 (1), 55–61. 10.1038/nrm3496.

(49) Reddy, G.; Straub, J. E.; Thirumalai, D. Influence of Preformed Asp23−Lys28 Salt Bridge on the Conformational Fluctuations of Monomers and Dimers of Aβ Peptides with Implications for Rates of Fibril Formation. J Phys Chem B 2009, 113 (4), 1162–1172. 10.1021/jp808914c.

(50) Itoh, S. G.; Yagi-Utsumi, M.; Kato, K.; Okumura, H. Key Residue for Aggregation of Amyloid-β Peptides. ACS Chem Neurosci 2022, 13 (22), 3139–3151. 10.1021/acschemneuro.2c00358.

(51) Khaled, M.; Rönnbäck, I.; Ilag, L. L.; Gräslund, A.; Strodel, B.; Österlund, N. A Hairpin Motif in the Amyloid-β Peptide Is Important for Formation of Disease-Related Oligomers. J Am Chem Soc 2023, 145 (33), 18340–18354. 10.1021/jacs.3c03980.

(52) Lopez del Amo, J. M.; Fink, U.; Dasari, M.; Grelle, G.; Wanker, E. E.; Bieschke, J.; Reif, B. Structural Properties of EGCG-Induced, Nontoxic Alzheimer’s Disease Aβ Oligomers. J Mol Biol 2012, 421 (4–5), 517–524. 10.1016/j.jmb.2012.01.013.

(53) Hsu, F.; Park, G.; Guo, Z. Key Residues for the Formation of Aβ42 Amyloid Fibrils. ACS Omega 2018, 3 (7), 8401–8407. 10.1021/acsomega.8b00887.

(54) Tomaselli, S.; Esposito, V.; Vangone, P.; van Nuland, N. A. J.; Bonvin, A. M. J. J.; Guerrini, R.; Tancredi, T.; Temussi, P. A.; Picone, D. The Α to β Conformational Transition of Alzheimer’s Aβ (1–42) Peptide in Aqueous Media Is Reversible: A Step by Step Conformational Analysis Suggests the Location of β Conformation Seeding. ChemBioChem 2006, 7 (2), 257–267. 10.1002/cbic.200500223.

(55) Wälti, M. A.; Ravotti, F.; Arai, H.; Glabe, C. G.; Wall, J. S.; Böckmann, A.; Güntert, P.; Meier, B. H.; Riek, R. Atomic-Resolution Structure of a Disease-Relevant Aβ(1–42) Amyloid Fibril. Proceedings of the National Academy of Sciences 2016, 113 (34). 10.1073/pnas.1600749113.

(56) Tycko, R. Solid-State NMR Studies of Amyloid Fibril Structure. Annu Rev Phys Chem 2011, 62, 279–299. 10.1146/annurev-physchem-032210-103539.

(57) Yang, Y.; Arseni, D.; Zhang, W.; Huang, M.; Lövestam, S.; Schweighauser, M.; Kotecha, A.; Murzin, A. G.; Peak-Chew, S. Y.; Macdonald, J.; Lavenir, I.; Garringer, H. J.; Gelpi, E.; Newell, K. L.; Kovacs, G. G.; Vidal, R.; Ghetti, B.; Ryskeldi-Falcon, B.; Scheres, S. H. W.; Goedert, M. Cryo-EM Structures of Amyloid-β 42 Filaments from Human Brains. Science (1979) 2022, 375 (6577), 167–172. 10.1126/science.abm7285.

(58) Liao, Q.; Owen, M. C.; Olubiyi, O. O.; Barz, B.; Strodel, B. Conformational Transitions of the Amyloid β Peptide Upon Copper(II) Binding and PH Changes. Isr J Chem 2017, 57 (7–8), 771–784. 10.1002/ijch.201600108.

(59) Ngo, S.; Guo, Z. Key Residues for the Oligomerization of Aβ42 Protein in Alzheimer’s Disease. Biochem Biophys Res Commun 2011, 414 (3), 512–516. 10.1016/j.bbrc.2011.09.097.

(60) Wu, K. Y.; Doan, D.; Medrano, M.; Chang, C. A. Modeling Structural Interconversion in Alzheimers’ Amyloid Beta Peptide with Classical and Intrinsically Disordered Protein Force Fields. J Biomol Struct Dyn 2022, 40 (20), 10005–10022. 10.1080/07391102.2021.1939163.

(61) Ciudad, S.; Puig, E.; Botzanowski, T.; Meigooni, M.; Arango, A. S.; Do, J.; Mayzel, M.; Bayoumi, M.; Chaignepain, S.; Maglia, G.; Cianferani, S.; Orekhov, V.; Tajkhorshid, E.; Bardiaux, B.; Carulla, N. Aβ(1-42) Tetramer and Octamer Structures Reveal Edge Conductivity Pores as a Mechanism for Membrane Damage. Nat Commun 2020, 11 (1), 3014. 10.1038/s41467-020-16566-1.

(62) Morimoto, A.; Irie, K.; Murakami, K.; Ohigashi, H.; Shindo, M.; Nagao, M.; Shimizu, T.; Shirasawa, T. Aggregation and Neurotoxicity of Mutant Amyloid β (Aβ) Peptides with Proline Replacement: Importance of Turn Formation at Positions 22 and 23. Biochem Biophys Res Commun 2002, 295 (2), 306–311. 10.1016/S0006-291X(02)00670-8.

(63) Tang, Z.; Chang, C. A. Systematic Dissociation Pathway Searches Guided by Principal Component Modes. J Chem Theory Comput 2017, 13 (5), 2230–2244. 10.1021/acs.jctc.6b01204.

(64) Maier, J. A.; Martinez, C.; Kasavajhala, K.; Wickstrom, L.; Hauser, K. E.; Simmerling, C. Ff14SB: Improving the Accuracy of Protein Side Chain and Backbone Parameters from Ff99SB. J Chem Theory Comput 2015, 11 (8), 3696–3713. 10.1021/acs.jctc.5b00255.

(65) Song, D.; Luo, R.; Chen, H.-F. The IDP-Specific Force Field *Ff14IDPSFF* Improves the Conformer Sampling of Intrinsically Disordered Proteins. J Chem Inf Model 2017, 57 (5), 1166–1178. 10.1021/acs.jcim.7b00135.

(66) Jorgensen, W. L.; Chandrasekhar, J.; Madura, J. D.; Impey, R. W.; Klein, M. L. Comparison of Simple Potential Functions for Simulating Liquid Water. J Chem Phys 1983, 79 (2), 926–935. 10.1063/1.445869.

(67) Essmann, U.; Perera, L.; Berkowitz, M. L.; Darden, T.; Lee, H.; Pedersen, L. G. A Smooth Particle Mesh Ewald Method. J Chem Phys 1995, 103 (19), 8577–8593. 10.1063/1.470117.

(68) Song, D.; Liu, H.; Luo, R.; Chen, H.-F. Environment-Specific Force Field for Intrinsically Disordered and Ordered Proteins. J Chem Inf Model 2020, 60 (4), 2257–2267. 10.1021/acs.jcim.0c00059.

(69) Mu, J.; Liu, H.; Zhang, J.; Luo, R.; Chen, H.-F. Recent Force Field Strategies for Intrinsically Disordered Proteins. J Chem Inf Model 2021, 61 (3), 1037–1047. 10.1021/acs.jcim.0c01175.

(70) Chen, J.; Fasihianifard, P.; Raz, A. A. P.; Hickey, B. L.; Moreno, J. L.; Chang, C.-E. A.; Hooley, R. J.; Zhong, W. Selective Recognition and Discrimination of Single Isomeric Changes in Peptide Strands with a Host: Guest Sensing Array. Chem Sci 2024, 15 (5), 1885–1893. 10.1039/D3SC06087J.

(71) Altis, A.; Nguyen, P. H.; Hegger, R.; Stock, G. Dihedral Angle Principal Component Analysis of Molecular Dynamics Simulations. J Chem Phys 2007, 126 (24). 10.1063/1.2746330.

(72) Mu, Y.; Nguyen, P. H.; Stock, G. Energy Landscape of a Small Peptide Revealed by Dihedral Angle Principal Component Analysis. *Proteins: Structure*, Function, and Bioinformatics 2005, 58 (1), 45–52. 10.1002/prot.20310.

(73) Chang, C.-E.; Potter, M. J.; Gilson, M. K. Calculation of Molecular Configuration Integrals. J Phys Chem B 2003, 107 (4), 1048–1055. 10.1021/jp027149c.

(74) Minh, D. D. L. Alchemical Grid Dock (AlGDock): Binding Free Energy Calculations between Flexible Ligands and Rigid Receptors. J Comput Chem 2020, 41 (7), 715–730. 10.1002/jcc.26036.

(75) Hikiri, S.; Yoshidome, T.; Ikeguchi, M. Computational Methods for Configurational Entropy Using Internal and Cartesian Coordinates. J Chem Theory Comput 2016, 12 (12), 5990–6000. 10.1021/acs.jctc.6b00563.

(76) Glorot, X.; Bengio, Y. Understanding the Difficulty of Training Deep Feedforward Neural Networks. In International Conference on Artificial Intelligence and Statistics; 2010.

(77) Kingma, D. P.; Ba, J. Adam: A Method for Stochastic Optimization. 2014.

(78) Michaud Agrawal, N.; Denning, E. J.; Woolf, T. B.; Beckstein, O. MDAnalysis: A Toolkit for the Analysis of Molecular Dynamics Simulations. J Comput Chem 2011, 32 (10), 2319–2327. 10.1002/jcc.21787.

(79) Paszke, A.; Gross, S.; Massa, F.; Lerer, A.; Bradbury, J.; Chanan, G.; Killeen, T.; Lin, Z.; Gimelshein, N.; Antiga, L.; Desmaison, A.; Köpf, A.; Yang, E.; DeVito, Z.; Raison, M.; Tejani, A.; Chilamkurthy, S.; Steiner, B.; Fang, L.; Bai, J.; Chintala, S. PyTorch: An Imperative Style, High-Performance Deep Learning Library. In Proceedings of the 33rd International Conference on Neural Information Processing Systems; Curran Associates Inc.: Red Hook, NY, USA, 2019.

