## Supporting Material for "Sampling Conformational Ensembles of Highly Dynamic Proteins via Generative Deep Learning"

**vBAT and BAT calculation details**

BAT coordinates are internal coordinate representation that is invariant/equivariant with respect to translations and rotations of molecular structure. In order to demonstrate how BAT and vBAT features are calculated, we use a simple four bonded atoms with indexes i, j, k, l whose positions are defined by $\vec{r}_{i}$, $\vec{r}_{j}$, $\vec{r}_{k}$, $\vec{r}_{l}$ (Figure S1).

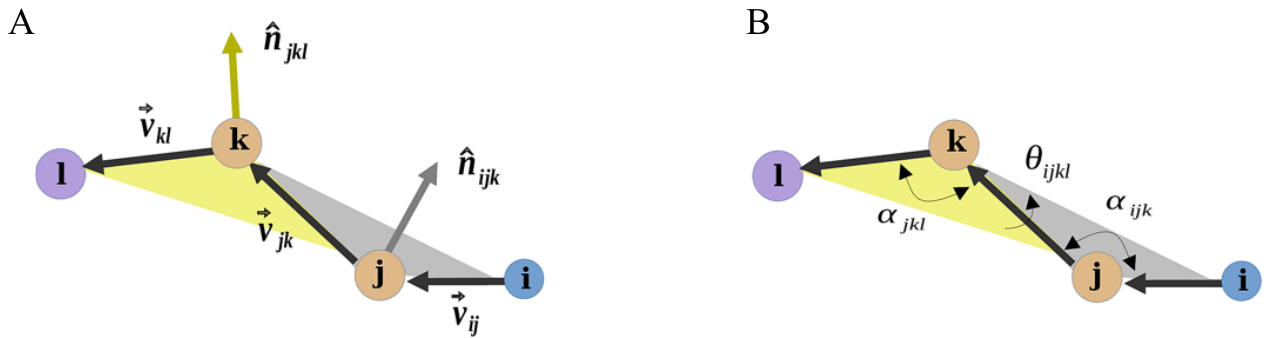

**Figure S1.** represents Vector BAT (vBAT) (a) and BAT (b) representations shown for a simple toy system, four bonded atoms with indexes i,j,k,l. In this, $\hat{n}_{ijk}$ is a unit vector normal to the gray plane determined by positions of atoms i, j, and k. $\hat{n}_{jkl}$ is another unit vector normal to the yellow plane determined by positions of atoms j, k, and l.

The bond vectors $\vec{v}_{ij}$ , $\vec{v}_{jk}$, $\vec{v}_{kl}$ that are the relative positions between the atoms, can be calculate as follows;

$\vec{v}_{ij}=\vec{r}_{i}-\vec{r}_{j}$, $\vec{v}_{jk}=\vec{r}_{j}-\vec{r}_{k}$, $\vec{v}_{kl}=\vec{r}_{k}-\vec{r}_{l}$. (1)

The unit vectors that are normal to the plane made by a pair of neighboring planar bond vectors define vector features, denoted as $\hat{n}_{ijk}$ and $\hat{n}_{jkl}$ .

$\hat{n}_{ijk}=\frac{-\vec{v}_{ij}\times\vec{v}_{jk}}{\left| \vec{v}_{ij} \right|\left| \vec{v}_{jk} \right|}$ , $\hat{n}_{jkl}=\frac{-\vec{v}_{jk}\times\vec{v}_{kl}}{\left| \vec{v}_{jk} \right|\left| \vec{v}_{kl} \right|}$	(2)

Equations 1 & 2 are essential for the calculation of bond length (b), bond angle (α) and dihedral angle (θ). For instance, the bond length between atoms i and j is the length of the vector $\vec{v_{ij}}$ etc.

$b_{ij}=\left| \vec{v}_{ij} \right|$ , $b_{jk}=\left| \vec{v}_{jk} \right|$ , $b_{kl}=\left| \vec{v}_{kl} \right|$	(3)

Bond angle between two vectors connected via central atom can be estimated via following equation;

$\alpha_{ijk}=arctan2\left( -\left| \vec{v}_{ij}\times\vec{v}_{jk} \right|,-\vec{v}_{ij}\cdot\vec{v}_{jk} \right)$ , $\alpha_{jkl}=arctan2\left( -\left| \vec{v}_{jk}\times\vec{v}_{kl} \right|,-\vec{v}_{jk}\cdot\vec{v}_{kl} \right)$. (4)

In this context, the cosine of the angle between planar vectors is determined by their dot product, while the sine of the angle is determined by their cross product. Solely relying on the cosine can introduce errors, particularly for angles spanning a wide range (-180, 180), such as torsion angles. To address this issue, we employ the *arctan2* function, which considers both the sine and cosine components. This approach ensures a more accurate transformation from vector space to angular space by placing the angle value in the correct quadrant.

$\theta_{ijkl}=arctan2\left( \frac{\left( \hat{n}_{ijk}\times\hat{n}_{jkl} \right)\cdot\vec{v}_{jk}}{\left| \vec{v}_{jk} \right|},\left( \hat{n}_{ijk}\cdot\hat{n}_{jkl} \right) \right)$	(5)

To convert BAT to Cartesian coordinates, the positions of three preceding atoms must be predetermined. Here, we assume that the positions of atoms j, k, and l are known, and we proceed to write down the equation to determine the position of atom i.

$\vec{r}_{i}=\vec{r}_{j}+b_{ij}\left( \vec{v}_{u}sin\left( \alpha_{ijk} \right)cos\left( \theta_{ijkl} \right)+\vec{v}_{p}sin\left( \alpha_{ijk} \right)sin\left( \theta_{ijkl} \right)-\vec{v}_{ij}cos\left( \alpha_{ijk} \right) \right)$	(6)

The new vectors $\vec{v}_{p}$ and $\vec{v}_{u}$ are calculated via following equations;

$\vec{v}_{p}=\frac{-\hat{n}_{jkl}}{\sqrt{1-\left( \frac{\vec{v}_{jk}\cdot\vec{v}_{kl}}{\left| \vec{v}_{jk} \right|\left| \vec{v}_{kl} \right|} \right)^{2}}}$, (7)

$\vec{v}_{u}=\frac{\vec{v}_{p}\times\vec{v}_{jk}}{\left| \vec{v}_{jk} \right|}$.	(8)

We have presented formulas for a simple four-atom system to enhance clarity and simplicity. However, it's worth noting that our GPU-optimized BAT code can efficiently perform these transformations for large systems, as demonstrated in the benchmark results illustrated in Figure S2.

**Evaluation of reconstructed conformations**

The accuracy of the ICoN model was assessed using the validation dataset. Initially, vBAT features extracted from the validation set were fed into the pretrained IcoN model, followed by a forward pass. Subsequently, the model's output was converted into fully atomistic Cartesian coordinates. Reconstruction accuracy was measured by computing the RMSD of all heavy atoms (including backbone atoms) between the original and reconstructed conformations. Heavy atom RMSD calculation involved all non-hydrogen atoms, while backbone RMSD estimation focused on N, Cα, C, and O atoms (Table 1).

Furthermore, we conducted more comprehensive comparisons between the original and reconstructed conformations, examining factors such as the distribution of all dihedral angles (including both backbone and sidechain for all amino acids) and their correlations. To address periodicity issues in angular space, dihedral angle correlations were computed using a trigonometric function (see details in the Supplementary Information) ^63,72,77^. These analyses were performed to verify that all properties of the original conformations are faithfully reproduced (see **Figure S3-S6**).

**Dihedral PCA (dPCA) calculation**

The dPCA calculation involves mapping torsion angles to a unit circle using trigonometric functions. This enables accurate estimation of differences (involving subtraction) and averages (involving summation) during the construction of the covariance matrix, thereby preventing erroneous computation of their correlation at the discontinuity margin (±180° or 360°/0°). For a dihedral angles θ and γ as variables of sample size N, covariance is calculated according to the following equation;

$Cov\left( \theta,\gamma\right)=\frac{1}{N}\sum_{i=1}^{n} \left( \theta_{i}-\overline{\theta} \right)\left( \gamma_{i}-\overline{\gamma} \right)$. (9)

The average value of the variable θ is estimated using following trigonometric trick;

$\overline{\theta}=arctan2\left( \frac{sin\left( \theta_{1} \right)+sin\left( \theta_{2} \right)+\ldots+sin\left( \theta_{n} \right)}{cos\left( \theta_{1} \right)+cos\left( \theta_{2} \right)+\ldots+cos\left( \theta_{n} \right)} \right)$ , (10)

while, subtraction is calculated using the following formula;

$\theta_{i}-\overline{\theta}=arctan2\left( \frac{sin\left( \theta_{i} \right)cos\left( \overline{\theta} \right)-sin\left( \overline{\theta} \right)cos\left( \theta_{i} \right)}{cos\left( \theta_{i} \right)cos\left( \overline{\theta} \right)+sin\left( \theta_{i} \right)sin\left( \overline{\theta} \right)} \right)$ . (10)

Using these formulas covariance can be calculated accurately avoiding errors due to periodicity in angular space.

**Generation of Novel Conformations**

*Step1: Non-linear Interpolation*

Non-linear interpolation is executed between consecutive points within a three-dimensional latent space. This process entails crafting a trajectory that transitions between the chosen point pairs $\vec{Z}_{i}$ and $\vec{Z}_{j}$, progressing along an arc while gradually altering the polar angle (Θ), azimuthal angle (Φ), and radius (R) within a spherical coordinate framework. Initially, a midpoint between the two points is determined by:

$\vec{Z}_{M}=\frac{\vec{Z}_{i}+\vec{Z}_{j}}{2}$ . (12)

Subsequently, we establish a positional vector for each point relative to the midpoint.

$\vec{R}_{i}=\vec{Z}_{i}-\vec{Z}_{M}$ , $\vec{R}_{j}=\vec{Z}_{j}-\vec{Z}_{M}$	(13)

Polar and azimuthal angles for both points are computed utilizing the following equations:

$\Theta_{i}=arccos\left( \frac{R_{i}^{x}}{\left| \vec{R}_{i} \right|} \right)$ , $\Phi_{i}=arctan2\left( \frac{R_{i}^{y}}{\left| \vec{R}_{i} \right|},\frac{R_{i}^{x}}{\left| \vec{R}_{i} \right|} \right)$ , (14)

$\Theta_{j}=arccos\left( \frac{R_{j}^{z}}{\left| \vec{R}_{j} \right|} \right)$ , $\Phi_{i}=arctan2\left( \frac{R_{i}^{y}}{\left| \vec{R}_{i} \right|},\frac{R_{i}^{x}}{\left| \vec{R}_{i} \right|} \right)$ . (15)

Subsequently, we determine an increment for each variable by dividing the interval between points i and j into N equal segments.

$\delta R=\frac{R_{j}-R_{i}}{N}$ , $\delta\Theta=\frac{\Theta_{j}-\Theta_{i}}{N}$ , ${dR}_{i}=\frac{R_{j}-R_{i}}{N}$ . (16)

To generate interpolation points, we systematically incorporate the increments into their respective variables as outlined below;

$R_{s}=R_{i}+s*\delta R$ , $\Theta_{s}=\Theta_{i}+s*\delta\Theta$ , $\Phi_{s}=\Phi_{i}+s*\delta\Phi$, (17)

where, ‘s’ represents the interpolation sample index ranging from 0 to N. A slight random adjustment is applied to the midpoint position before the commencement of each interpolation to generate a unique path. Upon completion of the interpolation process, all points are converted back to the Z space employing the following expression:

$\left( \begin{aligned} Z_{s}^{x} \\ Z_{s}^{y} \\ Z_{s}^{z} \end{aligned} \right)=\left( \begin{aligned} Z_{M}^{x} \\ Z_{M}^{y} \\ Z_{M}^{z} \end{aligned} \right)+R_{s}\left( \begin{aligned} cos\left( \Theta_{s} \right)sin\left( \Phi_{s} \right) \\ cos\left( \Theta_{s} \right)cos\left( \Phi_{s} \right) \\ cos\left( \Theta_{s} \right) \end{aligned} \right)$ . (18)

Equations 17 and 18 are iteratively applied across the entire sample set (N), yielding a non-linear trajectory that initiates at point i and transitions seamlessly to point j in a 3D latent space.

*Step 2: Eliminating non-physical conformations.*

To avoid steric clashes or non-physical conformations, all generated structures were subjected to a short minimization using a modified Generalized Born with implicit solvent model (mbondi3, igb=8) for 1000 steps. We applied energy cutoff of -400 (kcal/mol) and -200 (kcal/mol) for Aβ42 and aβ-crystallin57-69 respectively. All conformations with energy higher than the cutoff were excluded (**Table 1**). To have a fair comparison between various energy terms, the original MD trajectories were also minimized using a similar strategy involving 50 steps.

Step3: *Elimination of repeated conformations with pairwise RMSD search.*

After energy-based elimination, we further perform pairwise heavy atom RMSD search with 1Å cutoff across all remaining conformations in order to eliminate repeated conformations. 1 Å RMSD difference can capture small variations in sidechain as well as salt bridge formation or breaking. This step is essential to ensure all conformations are unique and diverse. **Table 1** shows the number of remaining conformations for all MD runs.

Step4: *Elimination of existing conformations in raw MD.*

We further compare remaining synthetic conformations from previous step with all the conformations in raw MD using pairwise heavy atom RMSD search. Conformations with the pairwise RMSD lower than 2 Å cutoff were deemed similar and excluded from synthetic pool for both Aβ42 and aβ-crystallin57-69 (**Table 1**). All pairwise RMSD searches were performed using PTRAJ ^78^.

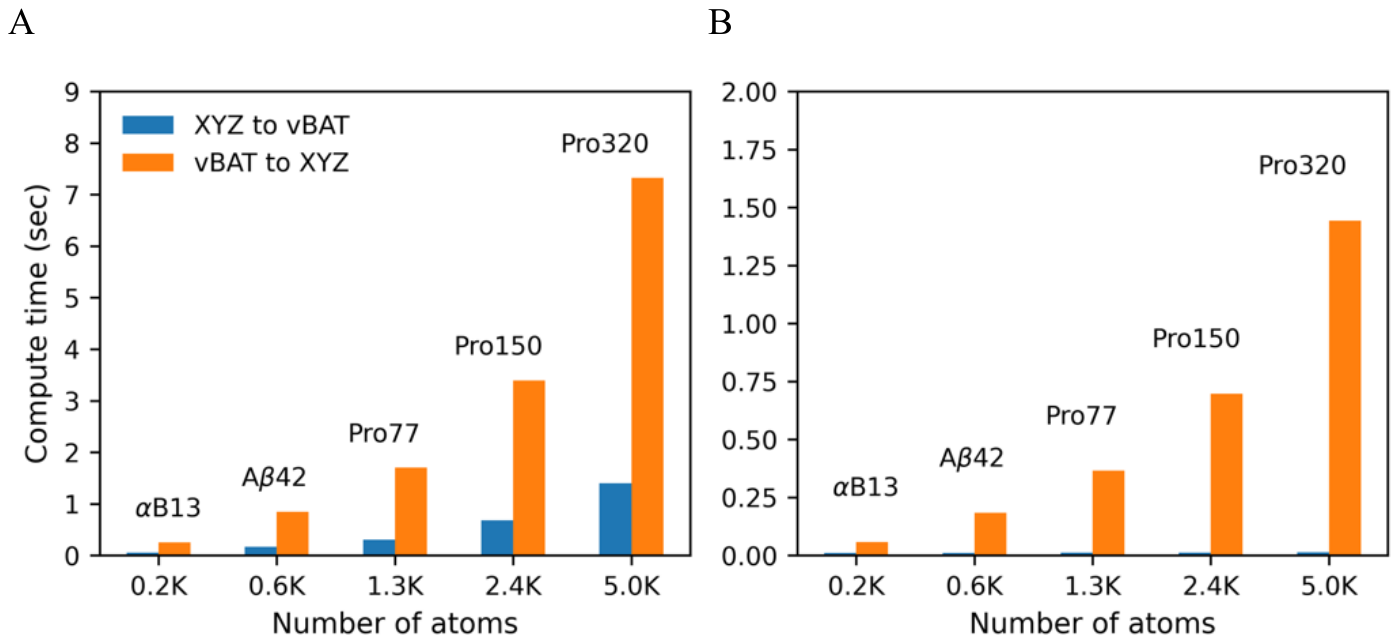
**Figure S2**. represents the benchmarks for both forward (Cartesian to vBAT) and backward (vBAT to Cartesian) transformations across proteins of various system sizes. The computation time for 5000 conformations for each system is reported. (A) shows the CPU performance, (B) illustrates the GPU performance. To fit the space constraints, the name 'αB-crystallin57-69' is abbreviated as 'αB13'.

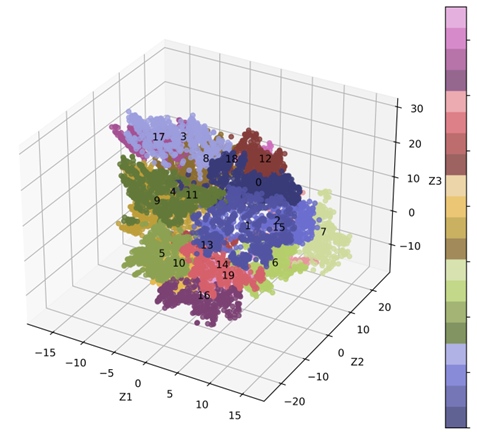

**Figure S3.** Latent space representation of 20 distinct clusters of αB-crystallin57-69.

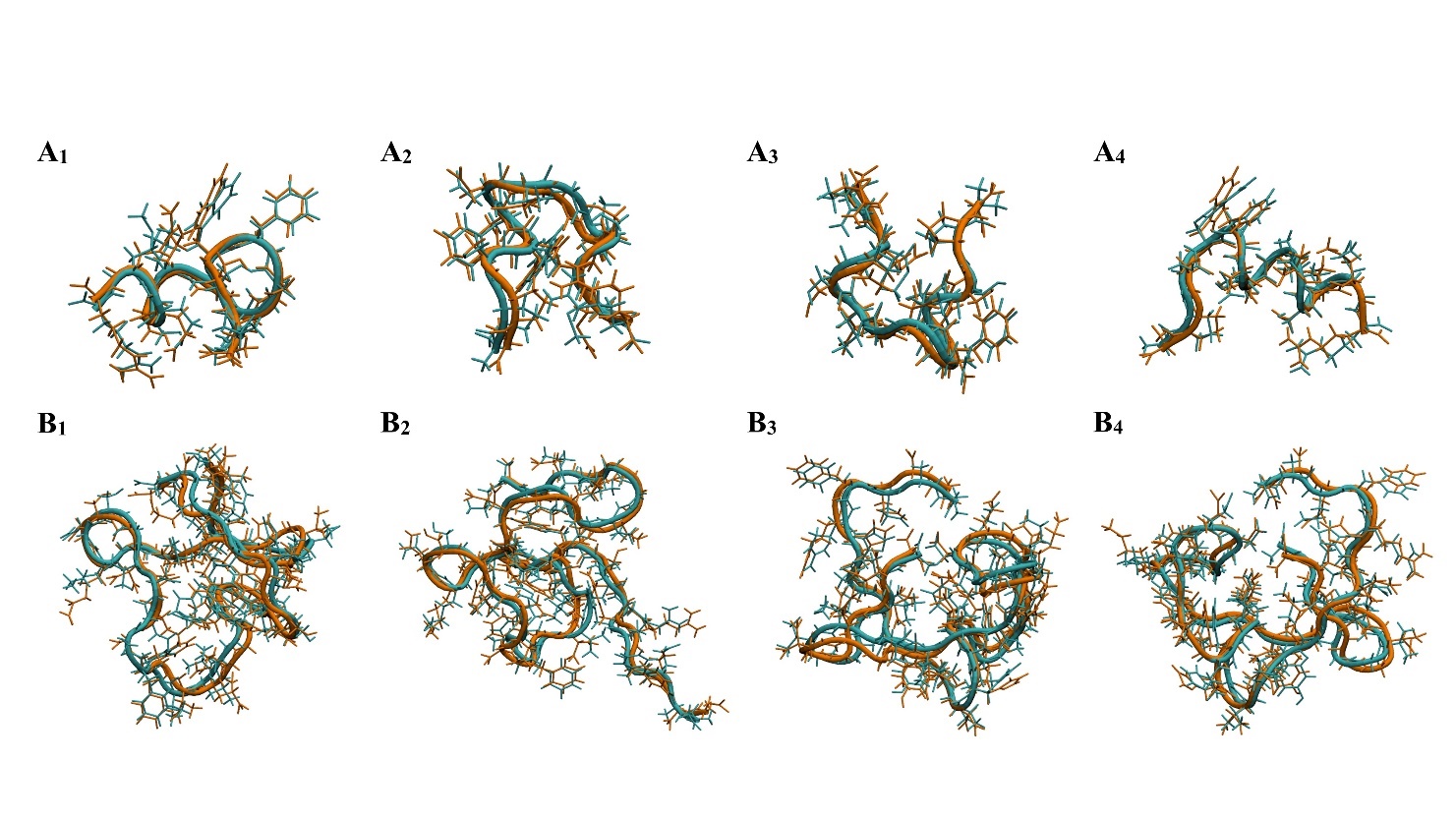
**Figure S4.** Comparisons of pairs of original and reconstructed conformations: four representative pairs of each system. The reconstructed (orange) and original (cyan) conformations all have heavy atom root mean square deviation (RMSD) within 0.9 Å for αB-cristallin57-69 (A1 to A4) and <1.3 Å for Aβ42 (B1 to B4).

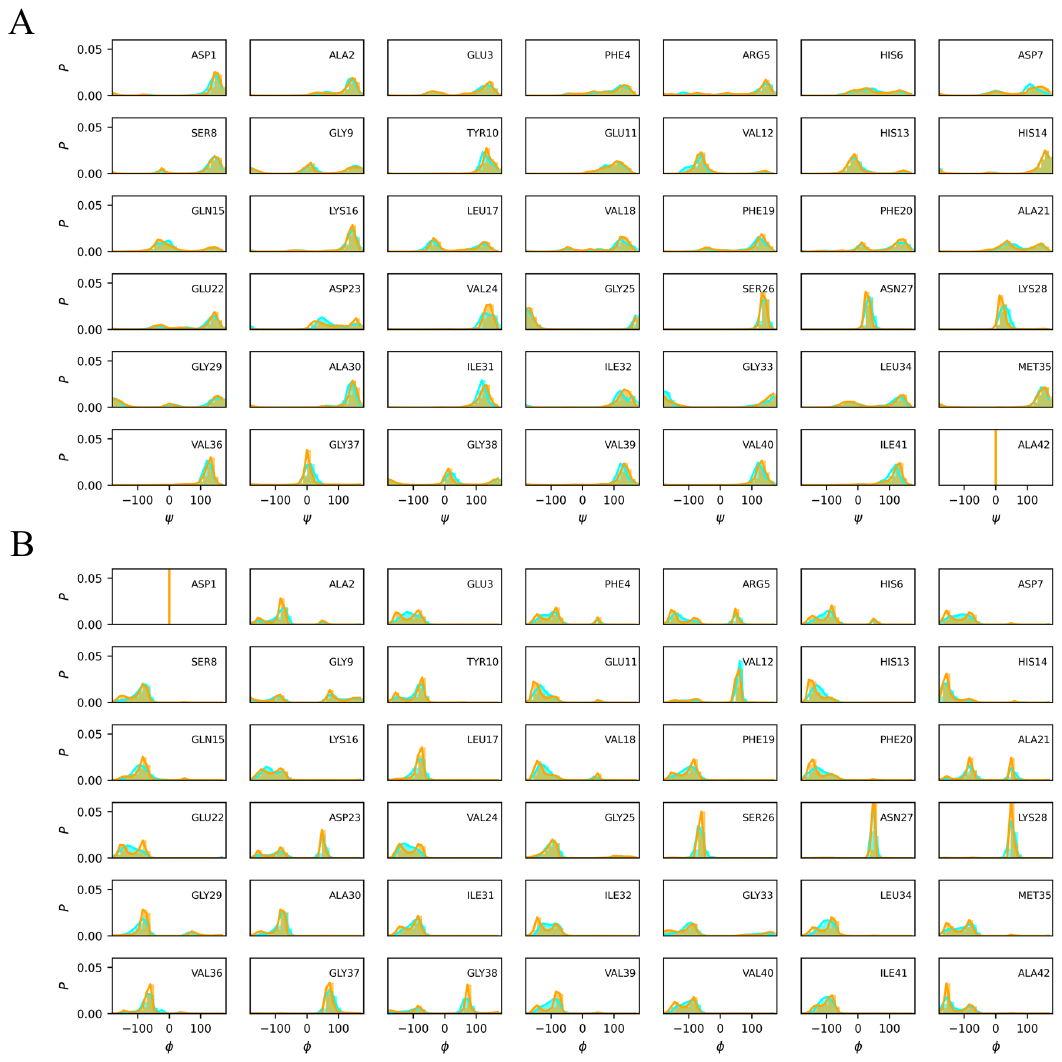

**Figure S5.** represents Aβ42 backbone dihedral angle (A) φ, and (B) ψ distributions shown for original MD conformations (cyan) and reconstructed validation set conformations (orange).

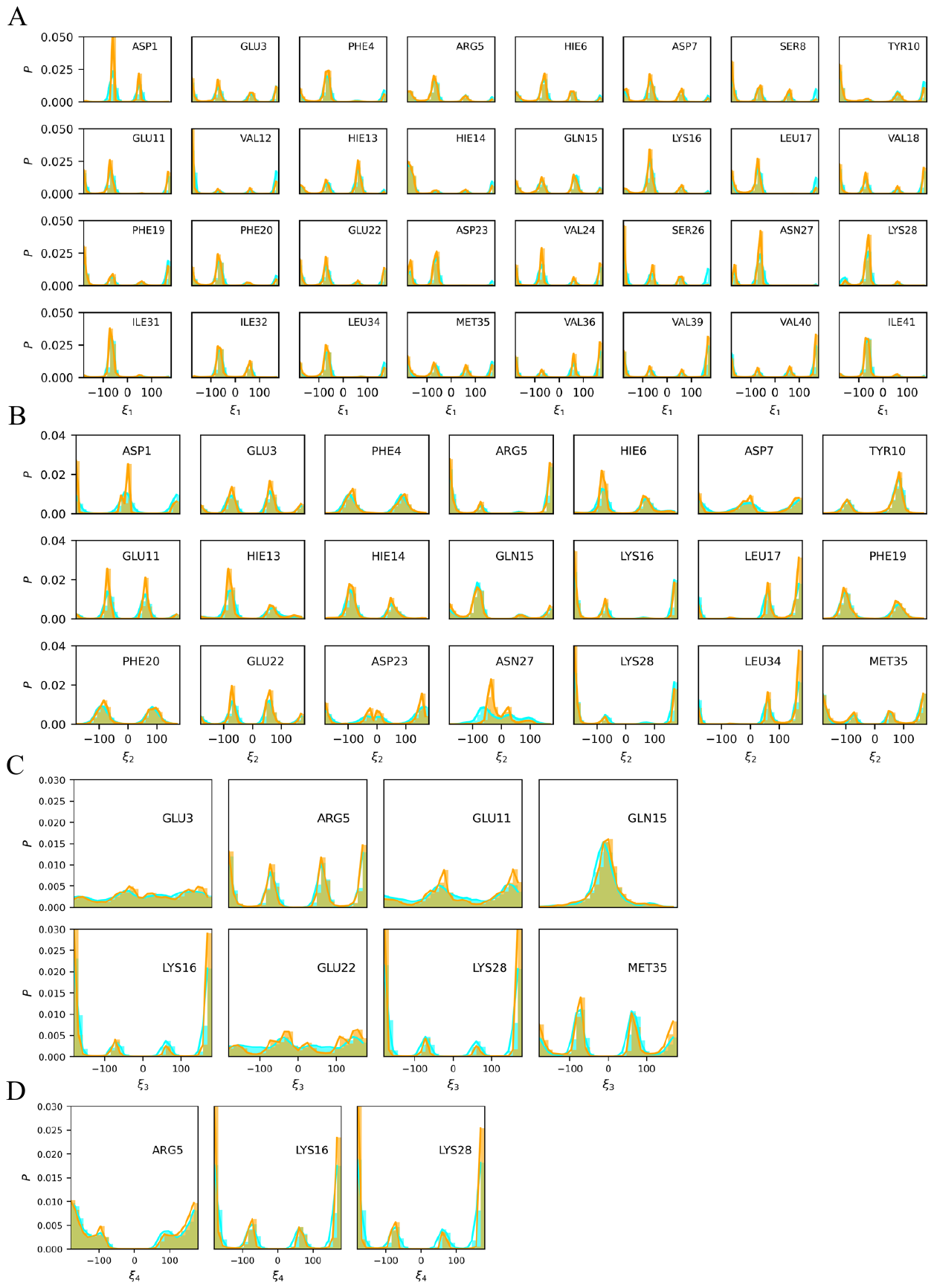

**Figure S6.** represents Aβ42 sidechain dihedral angle (A) χ_1_, (B) χ_2_, (C) χ_3_, and (D) χ_4_ distributions shown for original MD conformations (cyan) and reconstructed validation set conformations (orange).

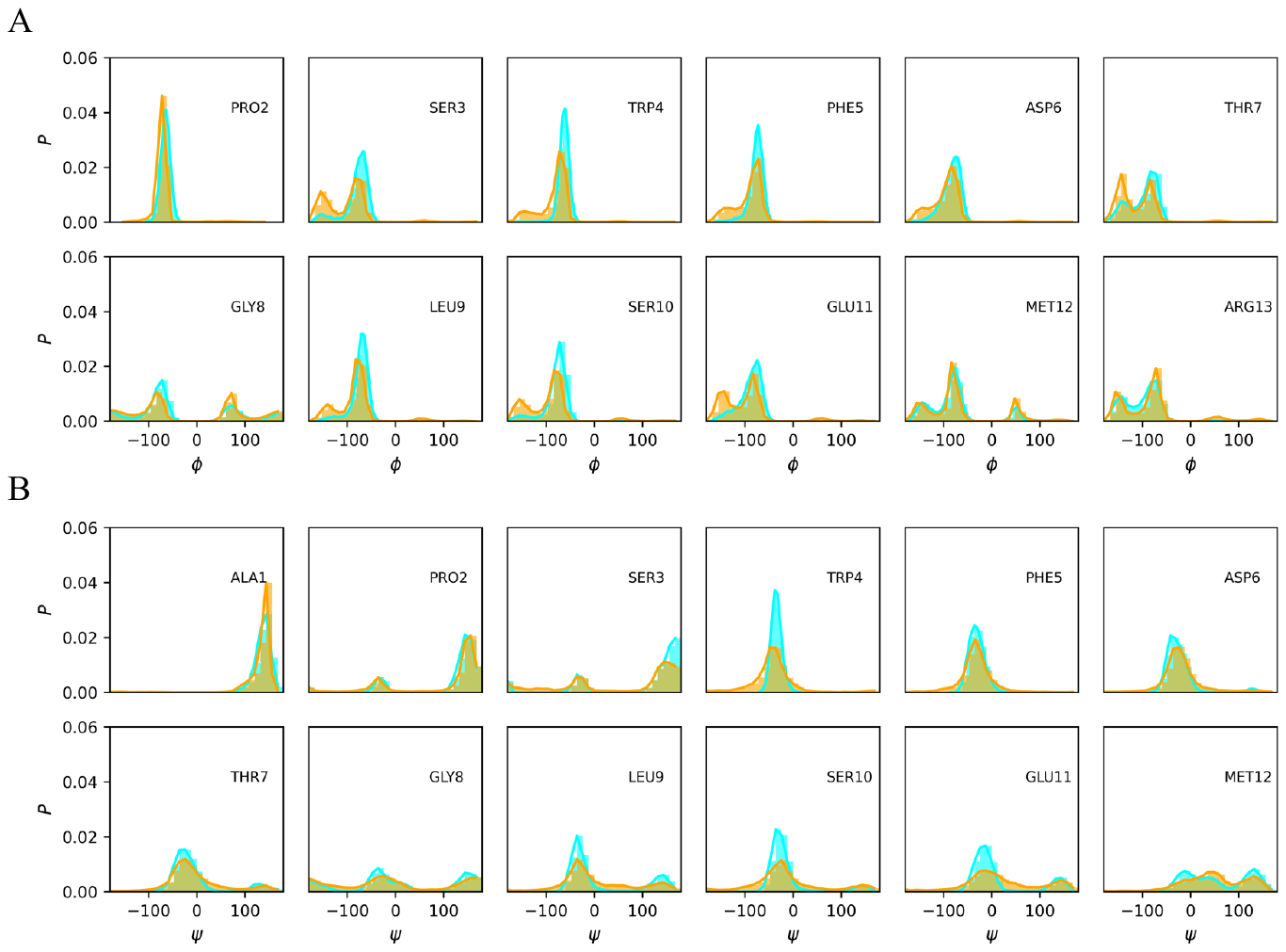
**Figure S7.** represents αB-crystallin57-69 backbone dihedral angle (A) φ and (B) ψ distributions shown for original MD conformations (cyan) and model generated reconstructed validation set conformations (orange).

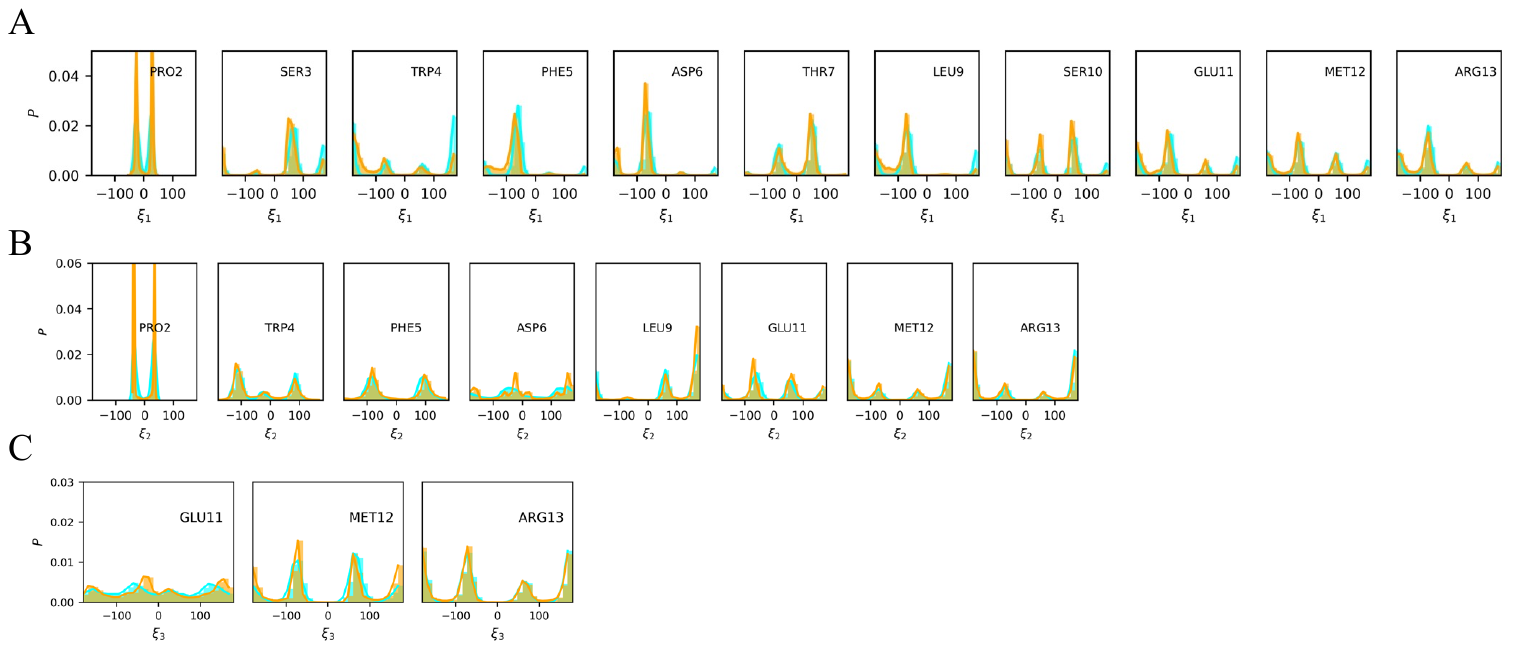
**Figure S8.** represents αB-crystallin57-69 sidechain dihedral angle (A) χ_1_, (B) χ_2_, (C) χ_3_ distributions shown for original MD conformations (cyan) and reconstructed validation set conformations (orange). The comparison shows that the ICoN model reconstructed conformations capture sidechain rotameric states accurately.

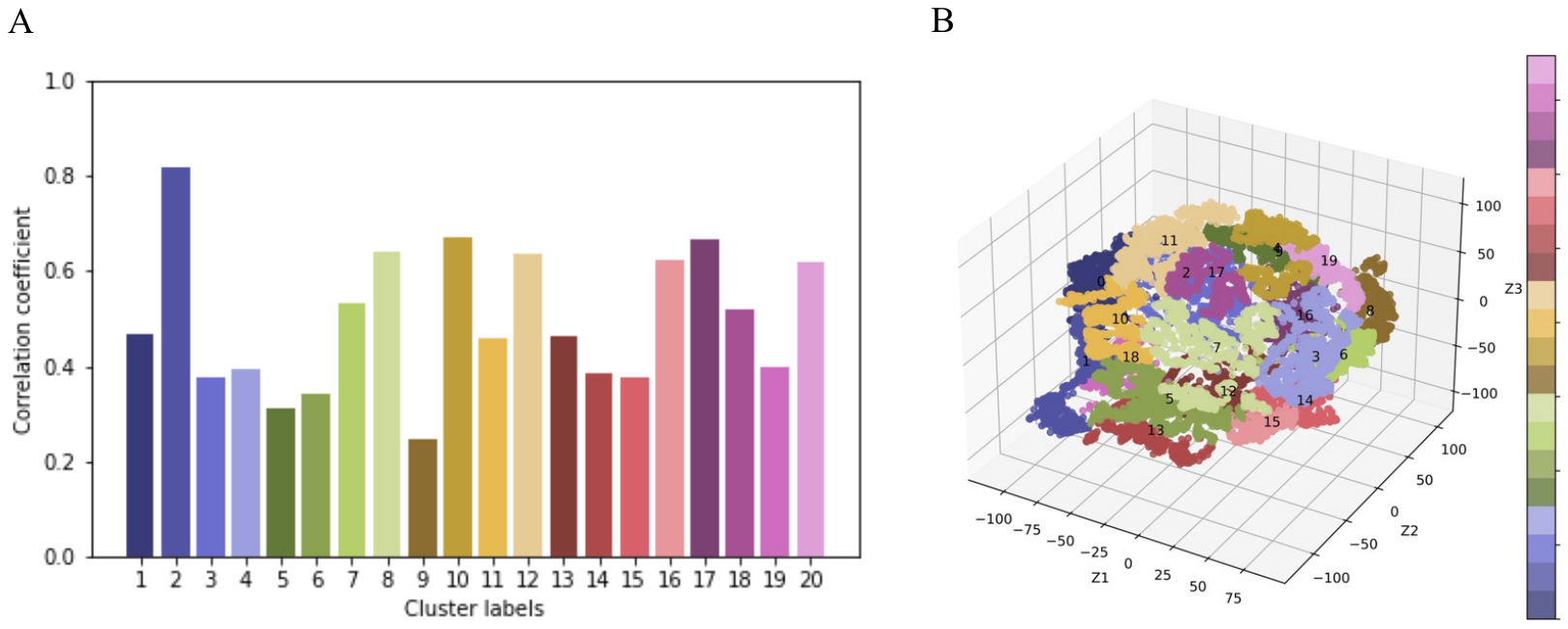

**Figure S9.** A point in 3D latent space can be transformed to a fully atomistic peptide structure, hence corresponding to a physical protein conformation. Structural similarity is quantified through pair RMSD within the conformational space, a measurement that can also be applied within the latent space. Consequently, we computed the correlations between pairwise distances within the latent space and pairwise RMSD values within the conformational space (A), across 20 distinct clusters (B) of Aβ42. The clustering process is conducted within the 3D latent space utilizing the Agglomerative Clustering method implemented through the *sklearn* library.

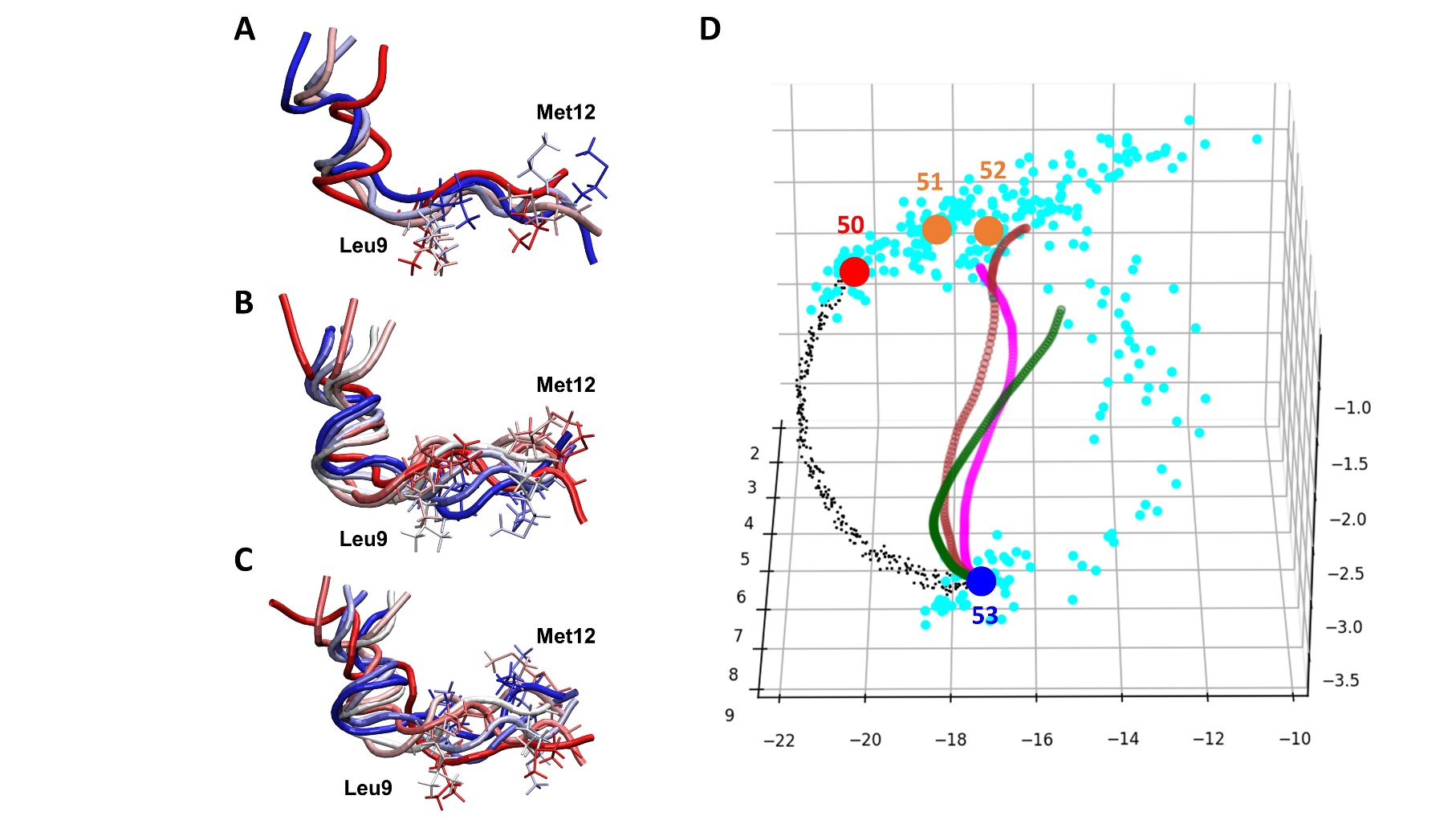
**Figure S10.** Conformational transitions of αB-crystallin57-69 peptide. Conformational transitions led by distortions using the (A) first dihedral principal component (PC) mode, (B) second PC mode and (C) first + second PC mode. Leu9 and Med12, which have the most fluctuations, are presented in licorice. Conformational transitions led by interpolation between training Conf indexes #50 (red dot) and #53 (dark blue dot). (D) 3D latent space from the ICoN model. Light blue dots present conformations saved every 1 ps in MD, with a total of 300 dots (300-ps simulation length). The MD frames were re-saved every 100 ps for training, and 4 frames are presented in the plot (red, blue and orange dots). The green dot line illustrates conformation distortions using the first PC mode, with the conformations shown in (A). The brown dot line illustrates conformation distortions using the second PC mode, with the conformations shown in (B). The magenta dot line illustrates conformation distortions using the first+second PC mode, with the conformations shown in (C).

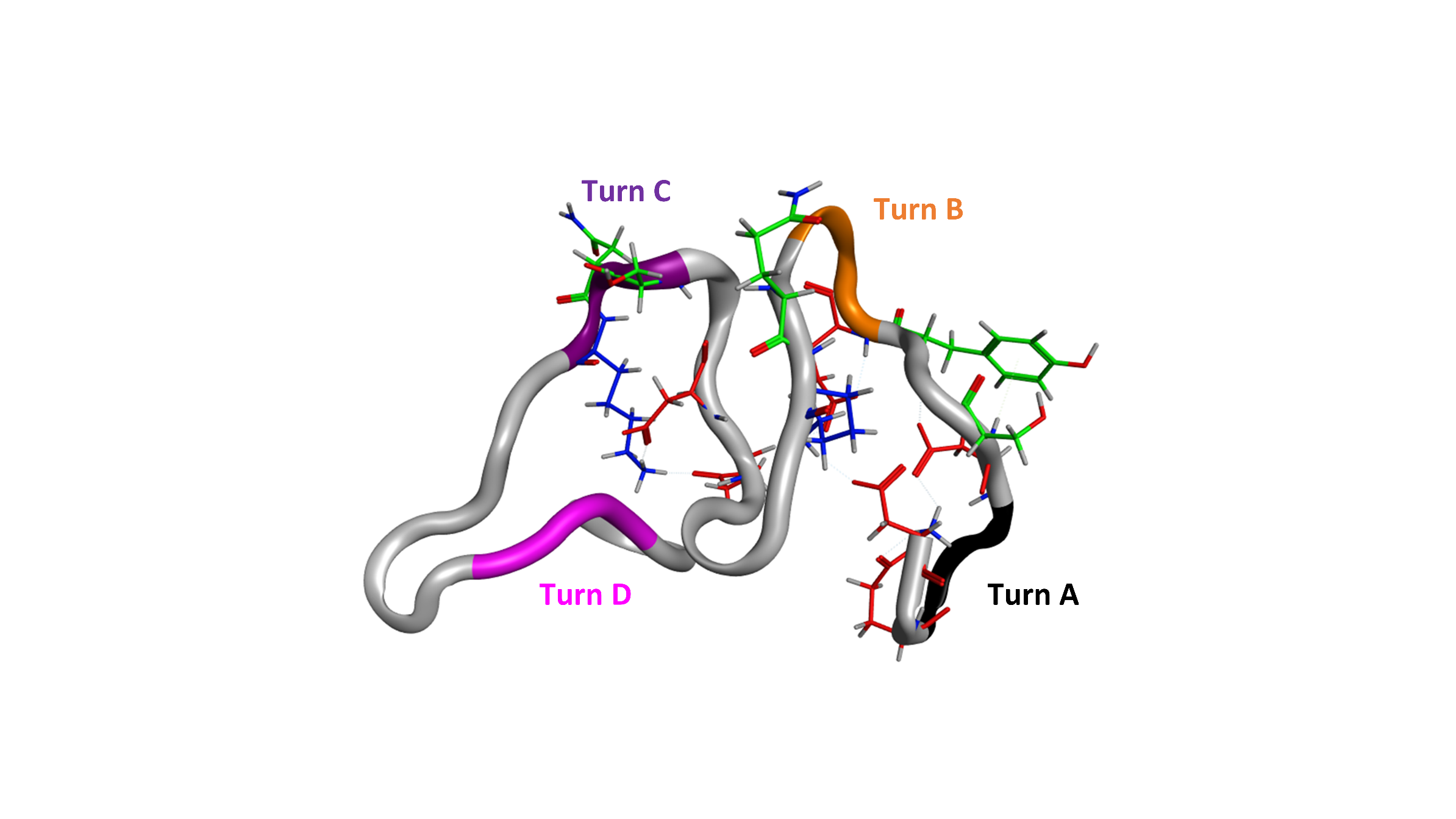

**Figure S11.** Initial conformations of MD run 5 with EFFS1 force field. Positively charged residue (blue). Negatively charged residue (red), and polar residue (green).

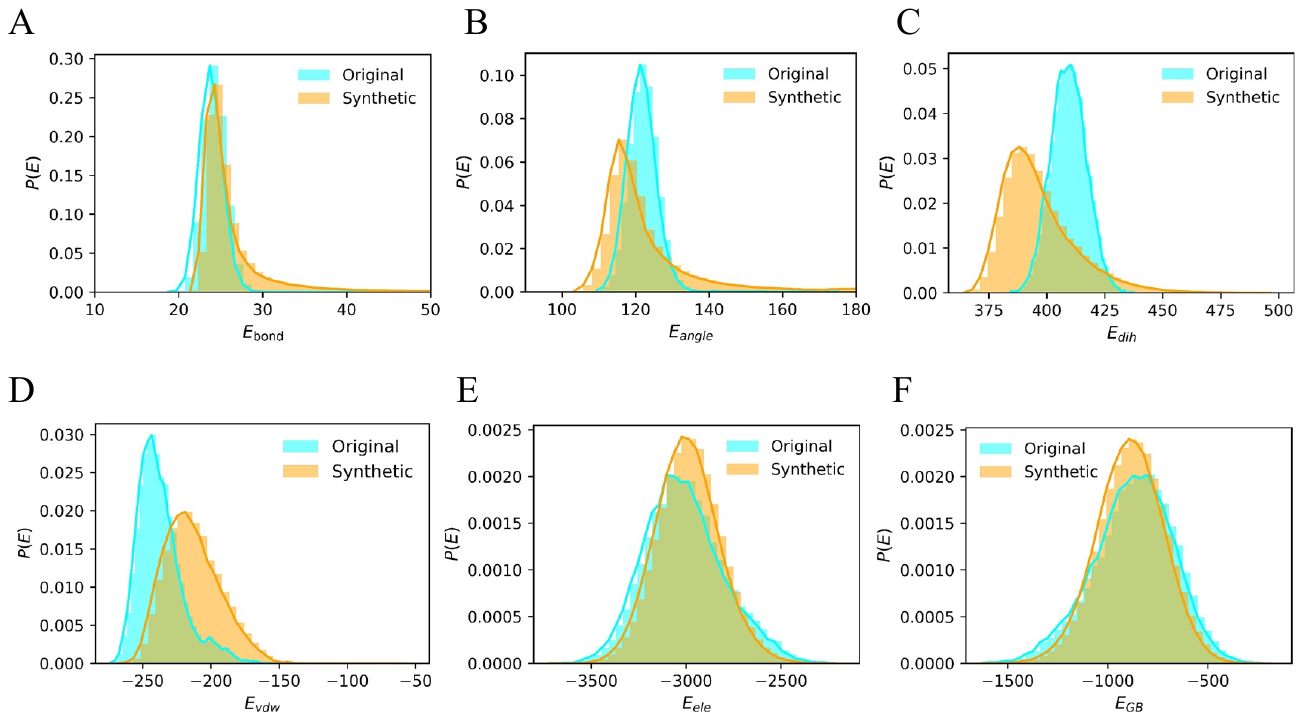

**Figure S12.** The energy distributions for different energy terms - (A) bond, (B) bond angle, (C) dihedral angle, (D) Van der Waals, (E) electrostatic interactions, and (F) Generalized Boltzmann (GB) (i) - are illustrated for the Aβ42 peptide. The energy distributions of MD and synthetically generated conformations by the ICoN model are shown in cyan and orange respectively. Distributions for synthetic conformations are calculated after energy-based elimination step, as explained in methods section. Comparison between MD and synthetic conformation energies emphasize that ICoN model is able to learn underlying physical properties of the system.

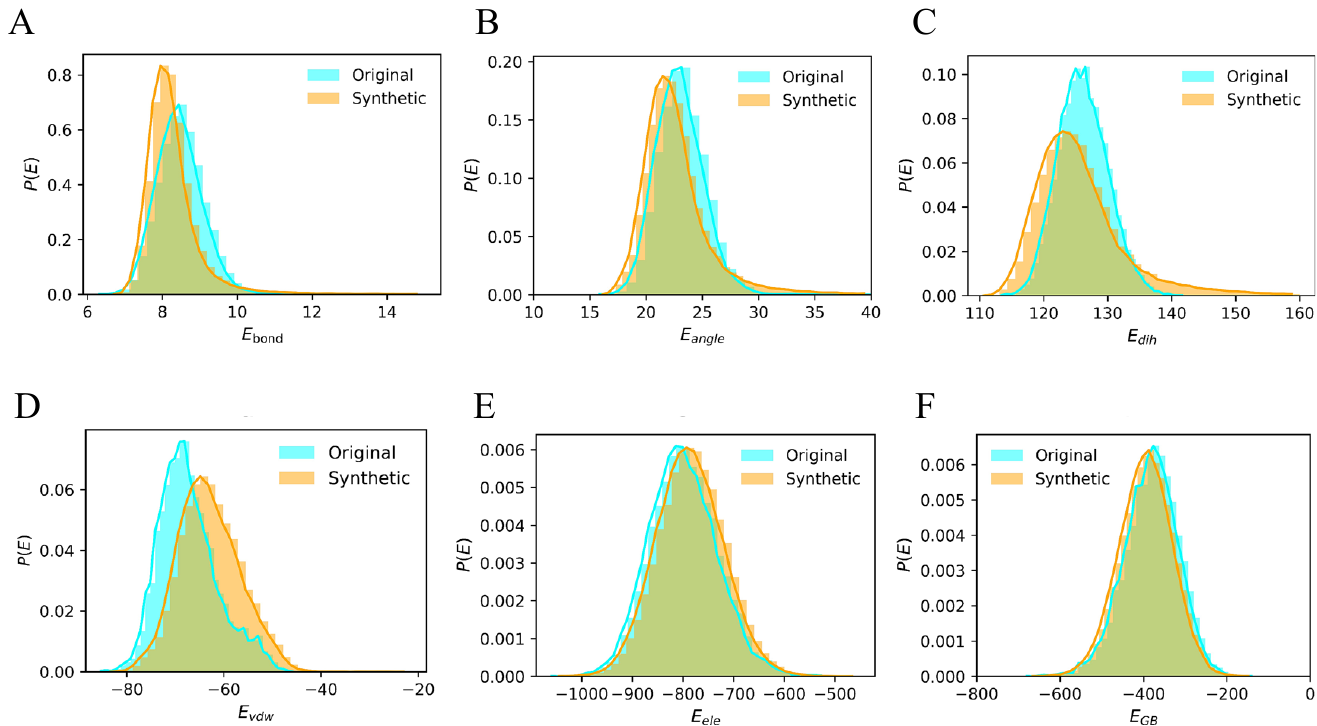

**Figure S13.** The energy distributions for different energy terms (A) bond, (B) bond angle, (C) dihedral angle, (D) Van der Waals, (E) electrostatic interactions, and (F) Generalized Boltzmann (GB) are illustrated for the αB-crystallin57-69 protein. The original molecular dynamics (MD) simulations' energy distributions are presented in cyan, while the energy distributions of synthetically generated conformations by the ICoN model are depicted in orange.

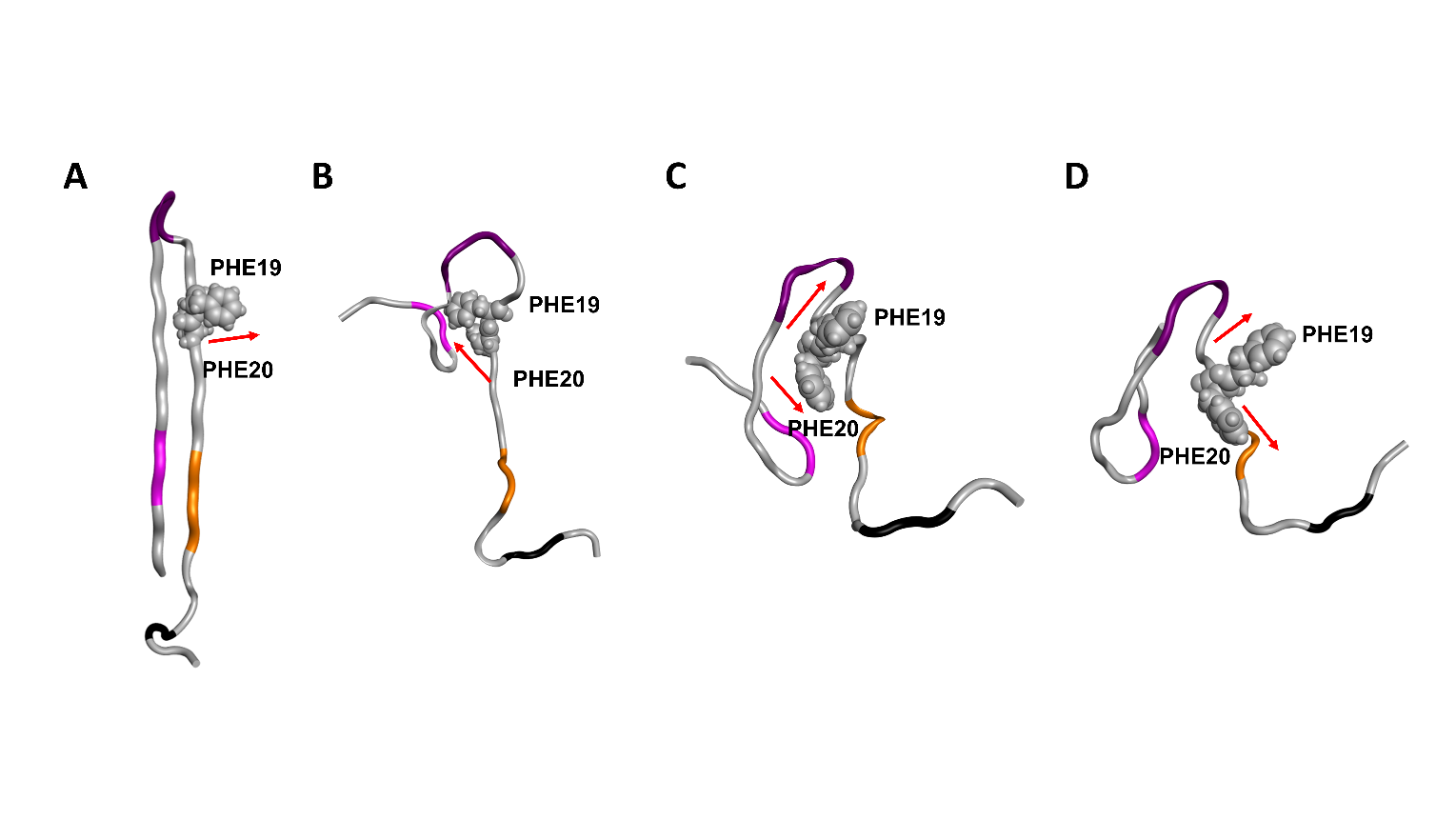
**Figure S14. Intermediate state with partial hair pin at Turn C conformations which require minimal conformational arrangement to form Aβ42 tetramer.** **(A)** Crystal structure of hair pin conformation that form Aβ42 tetramer (PDB: 6RHY). **(B-D)** Intermediate states with partial hair pin from novel synthetic conformations using FF14SB force field. Different orientations (direction indicated with red arrow) of F19 and F20 are observed leading to the formation of fibril conformation.

MD Run 5

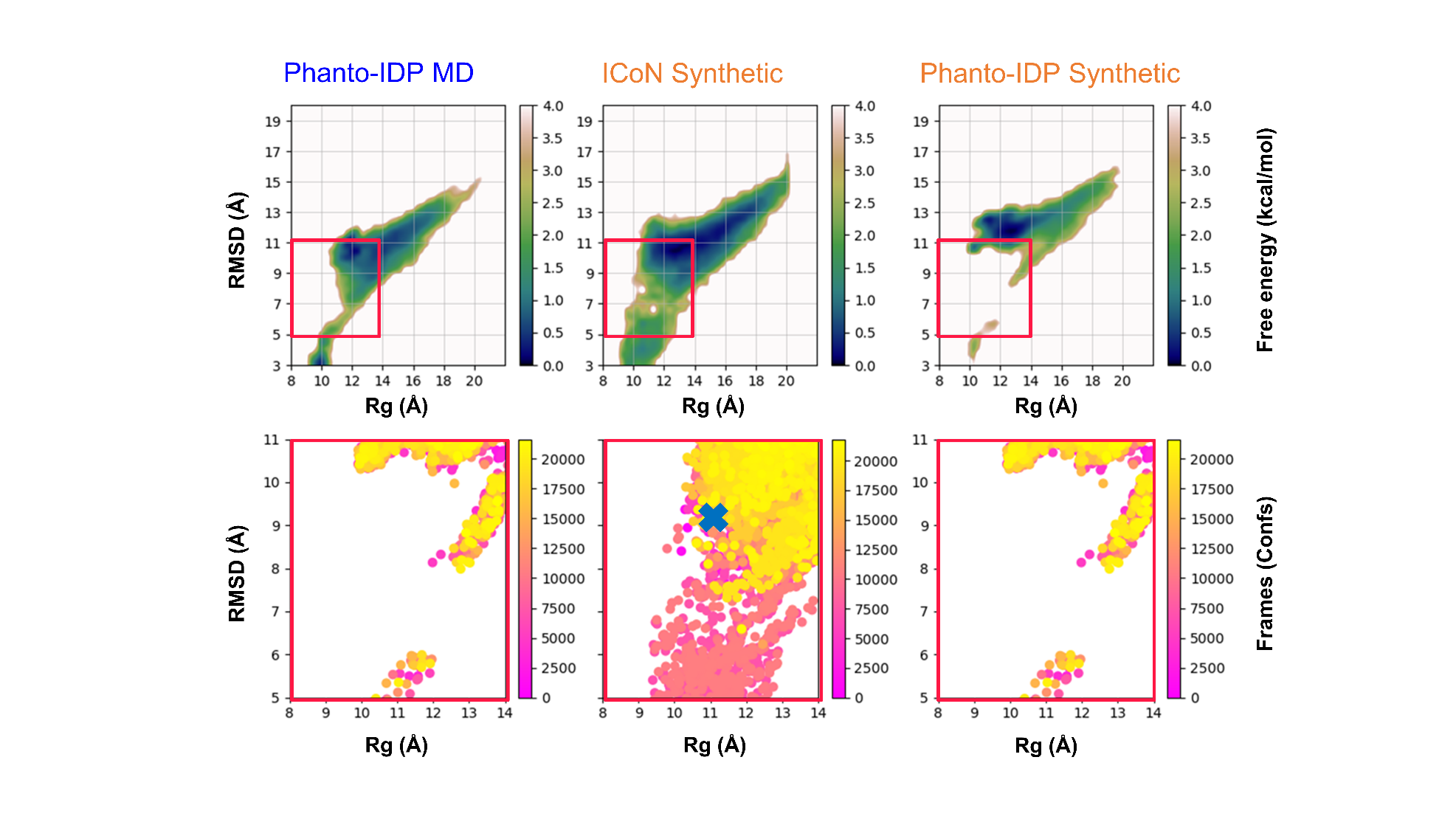

**Figure S15.** Comparing different machine learning models on the 2D space. Using MD Run 5 from Chen’s group as our training dataset, we applied the hyperparameters obtained from our MD Run 1 to sample new synthetic conformations with ICoN. For comparison, we added sidechains to backbone conformations generated by Phanto-IDP using AMBER tleap and carried out minimization and subsequent steps to remove repeat to obtain distinct newly synthetic conformations (see method for more detail). Red box indicates the transient state between initial and populated conformations. (**Top panel**) Free energy profile on the 2D space of RMSD and radius of gyration (Rg) for Aβ42. (**Bottom panel**) Zoom-in on the transient state (red box) of the top panel with color for each frame/conformation. The blue X indicates conformation select for Figure 7E.

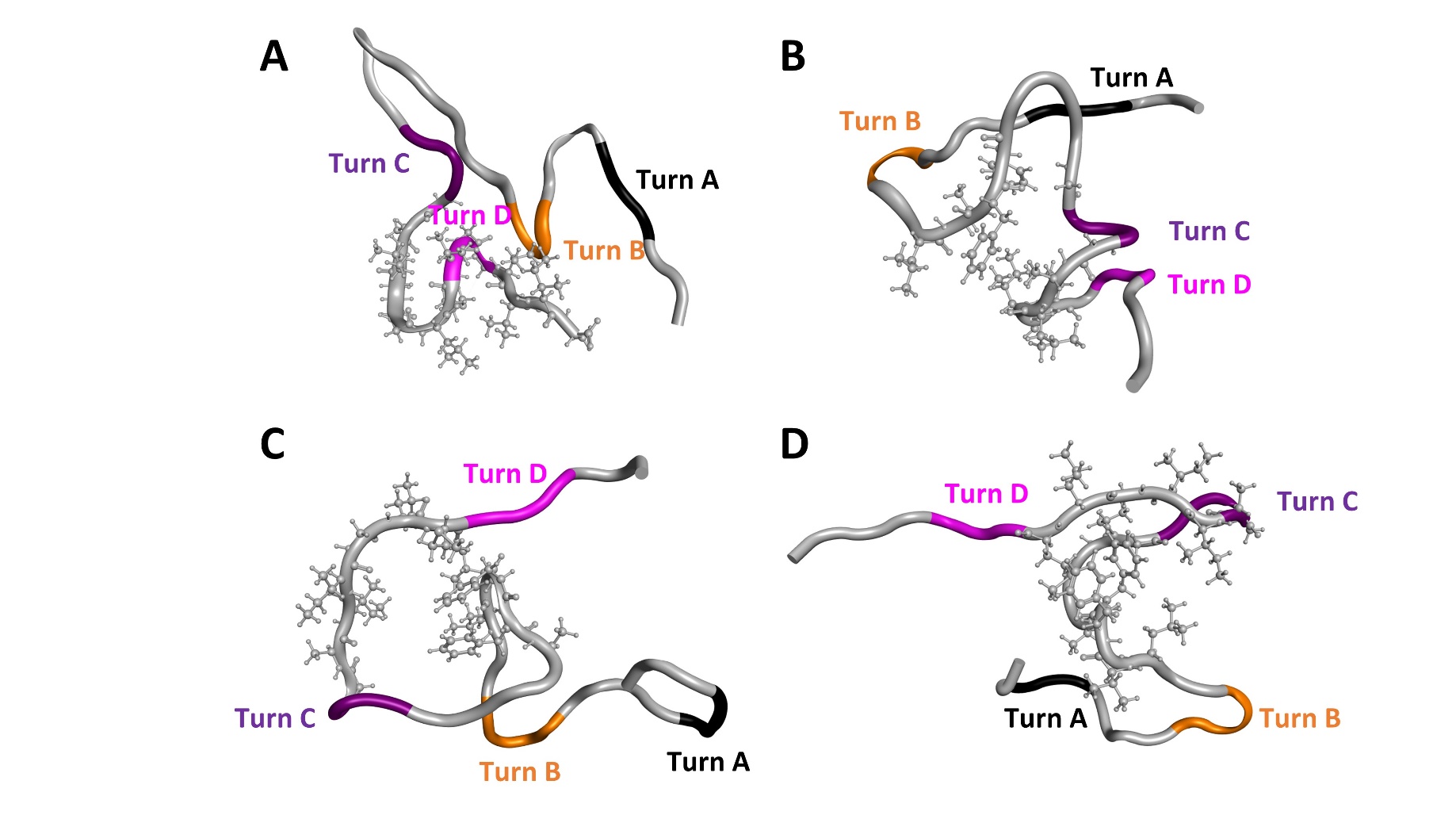
**Figure S16. Neurotoxic conformations of MD run5 with ESFF1 force filed in transient state** (red box in Figure S13). **(A)** Turns B, C and D presents with hydrophobic core around Turn D. **(B)** Turns B, C and D present with hydrophobic core around Turn C. **(C and D)** Turn A, B, C present with hydrophobic core around Turn C.

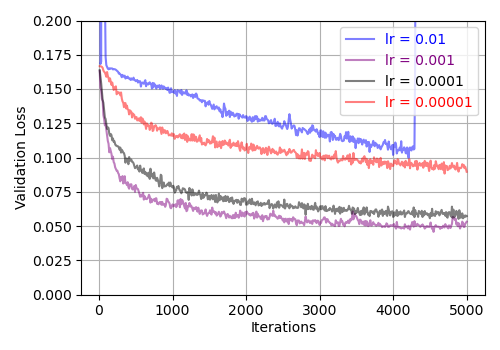

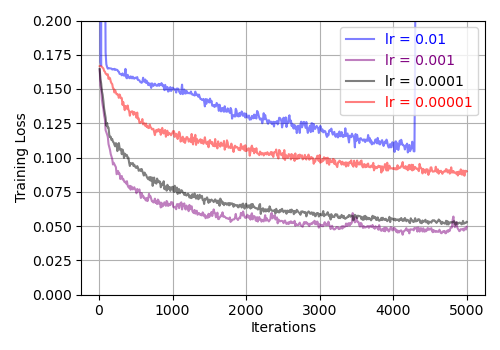
**Figure S17.** The learning curves for various learning rates were analyzed for both the training (Left) and validation (Right) sets of Aβ42 conformations. The curve with a learning rate of 0.00001 exhibited very slow convergence, whereas the curve at 0.01 was highly unstable. Learning rates of 0.001 and 0.0001 provided optimal and comparable performance. We selected a learning rate of 0.0001 due to its greater stability and convergence to a loss value similar to that achieved with 0.001.

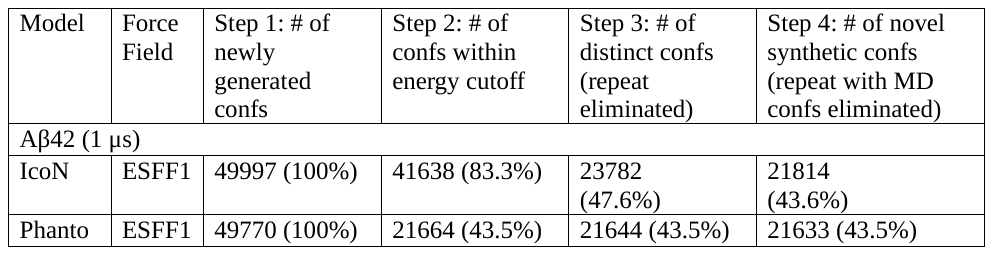
**Table S1.** Comparison of IcoN and Phanto models. Raw MD simulation and the novel synthetic conformations were obtained from Prof. Hei-Feng Chen’s group ^1^. Energy cutoff was 400 kcal/mol for Aβ42.Two conformations are treated as repeats when the computed heavy atom RMSD is smaller than 1 Å in Step 3. In Step 4, the RMSD cutoff is 1 Å.

**References:**

1.	Zhu, J. *et al.* Phanto-IDP: compact model for precise intrinsically disordered protein backbone generation and enhanced sampling. *Brief Bioinform* **25**, (2023).
